# DTX3L–PARP9 co-evolve as a single adaptive unit linking ubiquitination and ADP-ribosylation in antiviral immunity

**DOI:** 10.64898/2026.04.23.720386

**Authors:** Lea Picard, Emeline Esnouf, Thibault Latrille, Stéphanie Jacquet

## Abstract

Functional protein complexes are central to innate immunity, by integrating and diversifying signaling pathways of immune responses. Understanding how such complexes evolve is therefore key to elucidate the molecular basis of immune adaptation. Using a comparative genomics framework, we show that the DTX3L-PARP complex, which integrates ubiquitin-dependent and ADP-ribosylation pathways to regulate antiviral immunity, is an ancient and rapidly evolving functional module in vertebrates. We demonstrate that this complex has undergone recurrent adaptive evolution, with strong signatures of co-evolution between its components, consistent with early functional coupling and coordinated evolution as a single adaptive unit rather than independent proteins. Adaptive residues are enriched at protein–protein interaction surfaces within the RING-DTC domain of DTX3L, while PARP9 and PARP14 concentrate adaptive changes in regions involved in complex assembly and catalytic activities, consistent with hallmarks of pathogen-driven selection. These evolutionary signals remain detectable at short evolutionary timescales, including within species and sub-species, indicating ongoing adaptive evolution of the complex. These patterns support a model in which pathogens recurrently target the DTX3L–PARP axis to disrupt ubiquitin–ADP-ribose signaling, either by simultaneously interfering with catalytic domains or by destabilizing the complex interfaces. Altogether, our findings reveal the DTX3L–PARP9–PARP14 complex as a co-evolving adaptive module shaped by persistent host–virus arms races, highlighting how interconnected post-translational modification systems evolve in concert to sustain vertebrate antiviral immunity.

## INTRODUCTION

Protein complexes stand as fundamental evolutionary units in host innate immunity. By combining distinct signaling, enzymatic, and regulatory activities into functional assemblies, they expand the immune functional diversity, enabling new biological functions (Pereira-Leal et al., 2006) that drive rapid and coordinated responses to infection.

Post-translational modification (PTM) pathways are especially prone to such process, as they provide regulatory layers capable of shaping entire signaling pathways (Hsieh et al., 2026; Millán-Zambrano et al., 2022). By modulating protein activity, localization and interactions, PTMs contribute to the functional repertoire of eukaryote proteomes, and play a key role in the regulation of key cell processes, including innate immune responses. Among these modifications, ubiquitination and poly- or mono-ADP-ribosylation (PAR/MAR) are ancient and widely conserved across eukaryotes (Agrata & Komander, 2025; Palazzo et al., 2017). Although these systems originated independently, increasing evidence indicates that they became functionally interconnected during metazoan evolution (Barbour et al., 2023; Chatrin et al., 2025; Zhu et al., 2025), particularly in pathways involved in genome maintenance, stress responses, and innate immunity (Chatrin et al., 2025; Li et al., 2023). Ubiquitination involves the attachment of the small ubiquitin peptide to specific sites on target proteins (Agrata & Komander, 2025; Barbour et al., 2023), through the coordinated action of E1 (activation), E2 (conjugation), and E3 (ligase) enzymes. In parallel, ADP-ribosylation is catalyzed by members of the PARP family, which transfers ADP-ribose units from NAD⁺ onto target substrates as mono- or poly-ADP-ribose chains (Dasovich & Leung, 2023). Despite their mechanistic differences, they converge in multiple cellular contexts, particularly in pathways exposed to strong environmental pressures. This is especially evident in innate immunity, where ubiquitin-dependent modification contributes to pathogen sensing and clearance (Nag et al., 2023; Vozandychova et al., 2021), while ADP-ribosylation shapes antiviral signaling hubs (Du et al., 2023; Kar et al., 2024).

The DTX3L-PARP complex illustrates the evolutionary diversification and functional integration of these pathways. DTX3L is part of the DELTEX E3 ubiquitin ligases family which consists of five members (DTX1- DTX4 and DTX3L), all characterized by a C-terminal RING and an adjacent DELTEX C-terminal (DTC) domain (Scalia et al., 2023; Wang et al., 2021). Their N - terminal regions diverge: DTX1, 2 and 4 possess WWE domains, which are absent in DTX3 and DTX3L, suggesting functional specialization within the family. In vertebrates, DTX3L forms a functional complex with PARP9, in which PARP14 acts as a co-factor (Chatrin et al., 2020; Takeyama et al., 2003; C.-S. Yang et al., 2017), thereby linking ubiquitin-dependent and ADP-ribosylation pathways. The DTX3L-PARP9 complex plays central roles in DNA damage signaling and repair, transcriptional regulation, chromatin modulation, and carcinogenesis (Saleh et al., 2024; Yan et al., 2023; C.-S. Yang et al., 2017; Ye et al., 2024; Zhang et al., 2015). For example, it mediates mono-ubiquitination of histone H4 in response to DNA damage and regulates chromatin accessibility at interferon-stimulated genes through modification of histone H2B (Zhang et al., 2015). Recent work has further shown that DTX3L also binds and ubiquitinates ADP-ribose itself on single stranded nucleic acids (Zhu et al., 2024), further expanding this cross-modification network. As strong interferon-stimulated genes (ISGs), this complex also plays a key role in innate immunity (Kar et al., 2024; Zhang et al., 2015). Notably, DTX3L-PARP9 axis is critical for host defense against bacterial and viral infections (Huang et al., 2023; Thirunavukkarasu et al., 2023; Yan et al., 2023; Zhang et al., 2015), by modulating IFN production and signaling through STAT1 regulation, ubiquitinating viral 3C proteases for proteasomal degradation, and modifying host histone H2BJ to enhance antiviral state (Kar et al., 2024; Zhang et al., 2015). Moreover, the DTX3L–PARP9 complex regulates PARP14 activity through direct protein–protein interactions and post-translational mechanisms, thereby controlling interferon-γ–induced PARP14 ADP-ribosyltransferase activity (Kar et al., 2024; Saleh et al., 2024).

Because of its role in early viral sensing, interferon amplification, and antiviral response modulation, DTX3L-PARP complex represents a major candidate target of host–virus evolutionary dynamics. Several PARP family members, including PARP9 and PARP14, have been shown to exhibit signatures of adaptive selection in mammals, reminiscent of host-virus arm-races (Daugherty et al., 2014; Jacquet et al., 2025). In contrast, the evolutionary history of DTX3L remains poorly characterized, despite its central role in coordinating ubiquitin and ADP-ribose signaling. Given the physical and functional interactions among DTX3L, PARP9, and PARP14, we hypothesized that selective pressures may have acted on this complex as a whole functional unit. To address this, we analyzed the evolutionary history of DTX3L across vertebrates, and assessed the evolutionary forces shaping its diversification, using branch- and site-level analyses of positive selection with structural modeling of selected residues. We further integrated DTX3L into a phylogenomic comparative framework with PARP9 and PARP14 to evaluate signatures of co-evolution. We showed that DTX3L has experienced recurrent selective pressure since early metazoan diversification, with pronounced hallmarks in mammals. Importantly, we identified concordant patterns of selection across DTX3L, PARP9, and PARP14, consistent with co-evolution among the partners of this complex, that is likely driven by host–virus interactions. These dynamics are still detectable at the species and sub-species level, indicating recent and ongoing evolution. These findings uncover adaptive dynamics shaping a key ubiquitin–ADP-ribose regulatory axis in vertebrate immunity and illuminate how interacting post-translational modification systems can evolve in concert, highlighting the interplay between evolutionary constraint and adaptive diversification.

## RESULTS

### An ancient origin of DTX3L and early functional coupling with PARPs in vertebrates

To retrace the phylogenetic history of DTX3L, we retrieved orthologous sequences for all five DTX paralogs (Figure 1a) from 126 species spanning 21 metazoan orders and systematically assessed the genomic presence of the two PARP partners, PARP9 and PARP14 (Figure 1a, Supplementary Figure 1a). DTX-like sequences were found in distantly related lineages, although these could not be confidently assigned to specific paralogs, consistent with an early diversification followed by lineage-specific specialization (Figure 1b,c). Notably, we uncovered an ancient and evolutionary dynamic history of DTX3L, with an origin predating the radiation of extant Metazoa (Figure 1b,c). Phylogenetic reconstruction pointed to the DTX3L/DTX3 duplication as the founding event of the DELTEX gene family expansion, with other members of the family appearing through subsequent diversification events (Figure 1c). It also revealed a complex pattern of DTX3L gain and loss across metazoan evolution, with several clades (e.g., Insecta, Copepoda) exhibiting complete loss of all partners of the DTX3L-PARP9-PARP14 axis (Figure 1c). In contrast, following the divergence of Chondrichthyes, the presence of PARP9 correlated with the retention of the full DELTEX repertoire and the PARP partners, which suggests a shift toward DTX3L-PARP functional interdependence.

**Figure 1.**
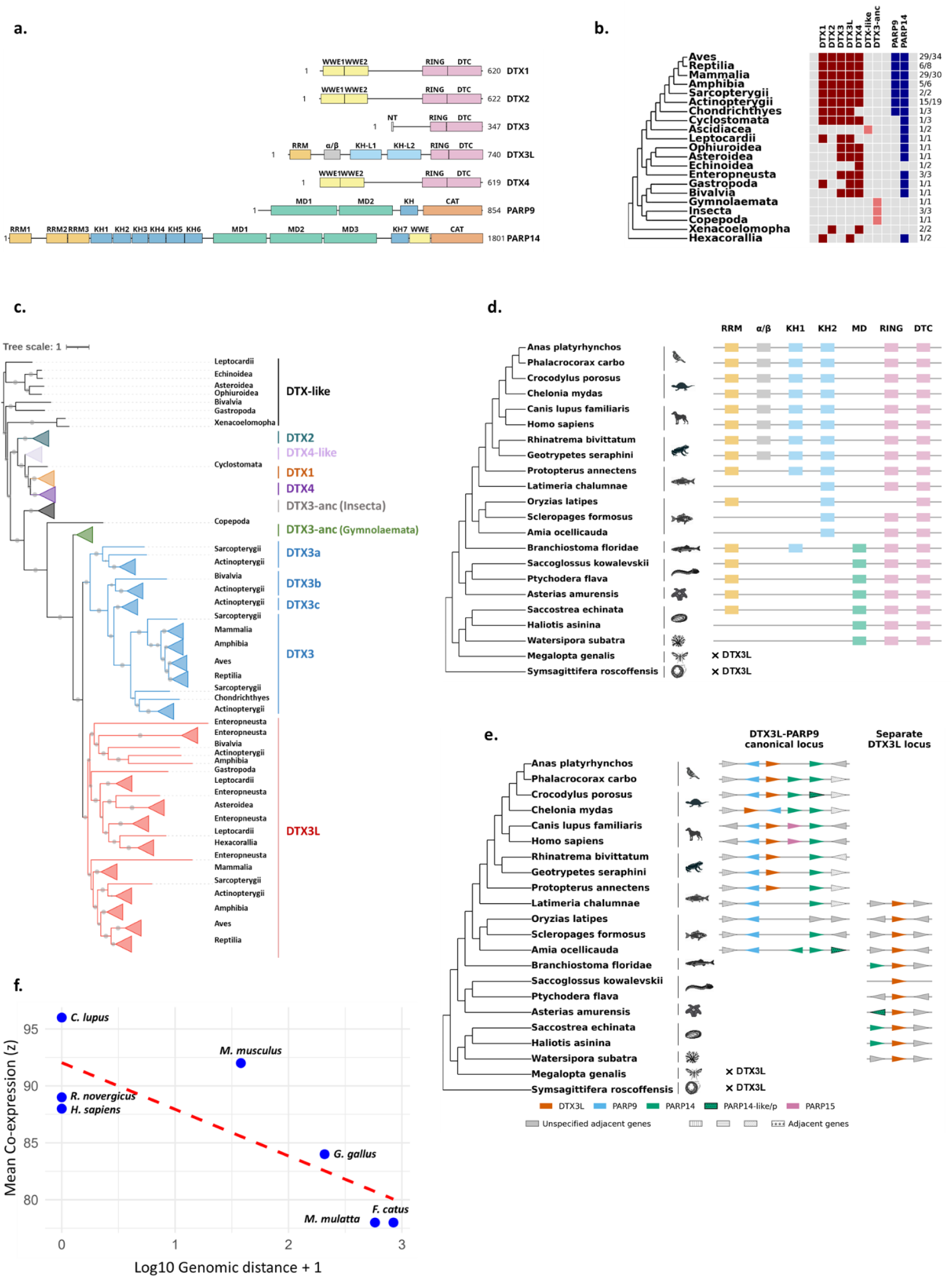
Ancient duplication of DTX3L and early functional coupling with PARP9. **a)** Domain architecture of Deltex family proteins, PARP9, and PARP14. Canonical domains are mapped along each protein using domain boundaries defined in InterPro, showing common and differing features between proteins from the same families. **b)** Phylogenetic representation of gene presence and absence across 21 metazoan classes. Filled cells indicate the presence of an ortholog, while empty cells denote absence. Deltex family proteins are shown in red (lighter red for inferred ancestral Deltex proteins) and PARP proteins in blue. Numbers on the right represent the number of species with the represented profile of presence and absence over the total number of species analyzed in the class. **c)** Maximum-likelihood phylogeny of the Deltex family reconstructed from full-length protein sequences. Major clades corresponding to DTX1–4 and DTX3L are labeled, with nodes support (bootstrap values) indicated by circles when > 0.7. **d)** Predicted domain architectures of DTX3L orthologs across representative metazoan species, illustrating conserved features and lineage-specific differences, and shared domains with PARP14. **e)** Graphical representation of the synteny of the canonical DTX3L locus across representative metazoan species and, the alternate PARP9-associated locus, showing the emergence of a conserved DTX3L-PARP9 locus following the diversification of dipnoi (lungfishes). **f)** Correlation between transcriptional expression (z) of DTX3L and PARP9 and their intergenic distance (Log10 genomic distance +1). Analysis was performed for different species (*Rattus norvegicus, Mus musculus, Homo sapiens, Macaca mulatta, Gallus gallus, Canis lupus* and *Felis catus*) show distinct intergenic distance, using available co-expression data from COXPRESdb. Significance of the slope was tested with Spearman test (ρ = −0.8, p-value = 0.029).

With few exceptions, most metazoan DTX3L orthologs possessed a RNA-Recognition Motif (RRM), along with the RING-DTC domain, indicating that RNA-binding capacity is an ancestral feature of the protein. The canonical structure of human DTX3L, comprising RRM, alpha/beta, 2 KH-L and RING-DTC domains, was conserved in the phylogeny as far back as amphibians (Figure 1d), although overall amino acid conservation varied among domains (Supplementary Figure 1b). In human, DTX3L shares a bidirectional UTR with PARP9 (Juszczynski et al., 2006), which likely promotes co-regulation and functional coupling between the two proteins. We therefore investigated the evolutionary origin and regulatory architecture of the DTX3L-PARP9 locus across metazoan species. We showed that DTX3L–PARP9 locus emerged after the diversification of Dipnoi (lungfishes), whereas the PARP9–PARP14 locus represents a more ancient genomic arrangement prior to fish diversification (Figure 1e). While the synteny between DTX3L and PARP9 was consistently maintained across tetrapod species, the organization of upstream neighboring genes varied between taxonomic orders, with changes in gene identity and orientation (Figure 1e), suggesting an evolutionary constraint on the physical coupling of DTX3L and PARP9. Interestingly, the emergence of PARP9 coincides with the loss of the macrodomain in DTX3L (Figure 1d, e), suggesting establishment of the functional dimer concomitant with PARP9 emergence, and take-over of the DTX3L macrodomain function by PARP9 macrodomains.

To further investigate the regulatory conservation across these proteins, we analyzed the intergenic region using the MultiZ alignment method implemented in UCSC Genome Browser (UCSC 100 vertebrates). Although synteny between DTX3L and PARP9 was broadly conserved among vertebrates, intergenic distance varied markedly across species, ranging from complete UTR sharing to several hundred nucleotides of distance, suggesting that UTR sharing is not an ancient, fixed genomic feature, but rather a lineage-specific arrangement, where genomic proximity may play a key role in functional coupling between the two genes. To assess whether intergenic distance influences transcriptional coordination, we extracted the Z-score ranking of the top 100 co-expressed genes for DTX3L, PARP9 and PARP14, from different species showing variable intergenic distance and available in the COXPRESdb database. We first tested the significance of co-expression between DTX3L, PARP9 and PARP14, then the correlation between co-expression strength and physical proximity. We demonstrated that the expression patterns of the three proteins significantly overlapped (p-value <10^-20^, Supplementary table 1), and that genomic distance was significantly and inversely correlated with co-expression strength (Spearman ρ = −0.8, p-value = 0.029), independent of phylogenetic relatedness (Figure 1f). Therefore, the closer the genes are, the higher the co-expression, supporting a model in which genomic proximity contributes to co-regulation. Interestingly, PARP14 consistently ranked among the top three genes co-expressing with DTX3L and PARP9 even in mammals which possess a PARP15 gene located upstream of DTX3L (Figure 1f, Supplementary Table 1). This transcriptional coupling was also significant between PARP9 and PARP14 in zebrafish, in which DTX3L is located in a distinct locus, indicating an early functional association among these partners in vertebrates.

### Extensive positive selection in DTX3L identifies RNA-binding and catalytic domains as candidate interfaces for viral antagonism

The strong evolutionary dynamics observed at the DTX3L locus suggest that it has been recurrently targeted by selective pressures throughout vertebrate evolution. We therefore applied the adaptive Branch-Site Random Effects Likelihood (aBSREL) (Smith et al., 2015) model to detect branches exhibiting accelerated rates of molecular evolution across four major vertebrate classes (mammals, birds, reptiles, and amphibians). We found strong signatures of positive selection at the ancestral nodes of each vertebrate class and along descendant branches, consistent with recurrent episodes of adaptive evolution (Supplementary Figure 2a). These episodes coincide with major phylogenetic divergence events in vertebrates, suggesting that DTX3L experienced adaptive changes during key evolutionary transitions. As positive selection was particularly pronounced in reptiles and mammals (Supplementary Figure 2a), we therefore examined the patterns of episodic diversification across representative mammalian orders, for which high-quality representative genomes are available, including primates, rodents, artiodactyls and carnivores. We showed that DTX3L has experienced differential selective regimes and lineage-specific adaptive pressures, with bats and rodents exhibiting the greatest number of branches under significant positive selection (Figure 2a). Given the episodic signatures of positive selection detected in DTX3L across mammals, we next investigated these signals at gene-wide and codon-specific levels. Both types of analyses, gene-level using the Branch-site Unrestricted Statistical Test for Episodic Diversification (BUSTED) (Murrell et al., 2015) and site-level inference implemented in Bio++ (Guéguen et al., 2013) and Codeml (PAML) (Z. Yang, 2007), showed strong evidence for ancient and recurrent diversifying selection in all five orders (p-value < 0.005), in contrast to the other DTX members, except DTX3 (Supplementary Figure 2b). A high proportion of codons were detected under significant positive selection ranging from 5.40 to 11.05% across lineages, with the highest percentages observed in bats, primates, and artiodactyls (Figure 2b), consistent with intensified selective pressures in these clades.

**Figure 2.**
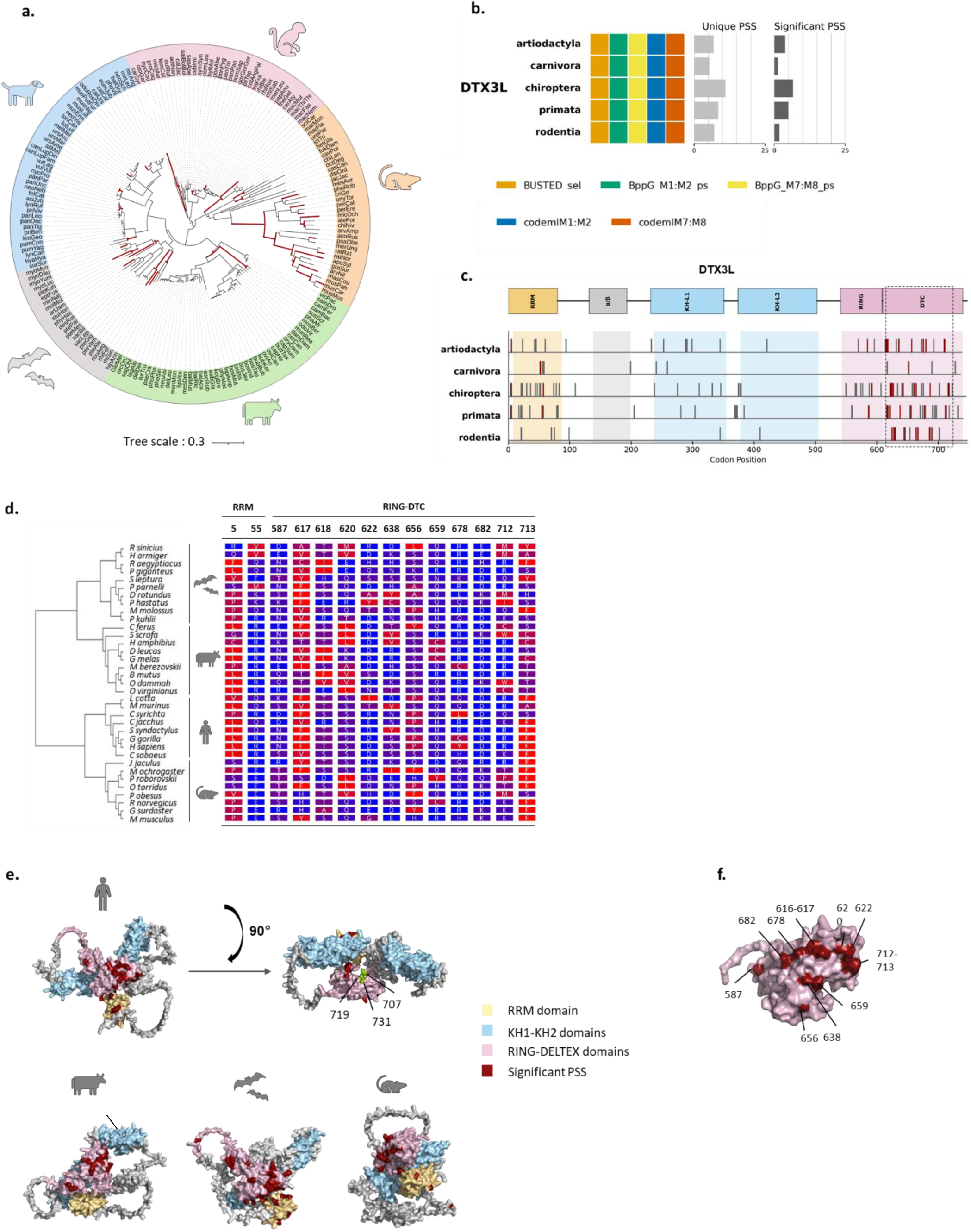
Evolutionary hotspots reveal adaptive shared substitutions in catalytic domains of mammalian DTX3L. **a)** Maximum likelihood phylogenetic trees of mammalian (primates, rodents, artiodactyls, bats and carnivores) DTX3L, showing branches under significant positive selection (p-value <0.05, in red). **b)** Summary of BUSTED (Branch-site Unrestricted Statistical Test for Episodic Diversification) and models M0, M1, M2, M7, and M8 implemented in Bio++ and codeml (PAML) with nested model comparisons (M1 vs. M2 and M7 vs. M8) assessed by likelihood ratio tests (LRTs) on DTX3L across mammalian orders. Colored boxes indicate models detecting a significant signal of positive selection and barplots indicate the proportion of all positively selected sites (Unique PSS) and Significant PSS (shared by at least two models). **c)** Significant positively selected sites mapped on the linear representation of DTX3L protein structure for each mammalian order. Red bars indicated the PSS shared by at least three mammalian orders. The dotted line box show the stretch of shared PSS. Codon numbering is based on the reference sequences for each order: artiodactyla - *Bos mutus*, carnivora - *Canis lupus*, chiroptera - *Pteropus giganteus*, primata - *Homo sapiens*, rodentia - *Mus musculus*)**. d)** Alignments of positively selected sites shared by at least three mammalian orders, highlighting amino acid changes in mammalian DTX3L. The cladogram on the left shows the phylogenetic relationships among mammalian species. **e)** Predicted three-dimensional structures of DTX3L generated using AlphaFold2 for three representative mammalian species*: H. sapiens, B. mutus*, and *P. giganteus*. **f)** Close-up view of the DTC domain of DTX3L, where shared significant PSS across at least three orders are shown in red.

Although adaptive substitutions were widespread along the protein, most of them clustered preferentially and significantly within canonical functional domains, notably in the RNA recognition motifs (RRM) and the catalytic RING and DTC domains (Kruskal-Wallis test, 10^-8^ < p-value < 0.02; pairwise Wilcoxon tests, p-value < 0.05) (Figure 2c, Supplementary Table 2). These patterns were similar across mammals, suggesting that adaptive pressures have continually acted on regions that are central to RNA binding and ubiquitin ligase activity. Interestingly, a cluster of 10-15 rapidly evolving codons was identified within the DTC domain (positions 616–697 in human, Figure 2c). This pattern was systematically observed across all mammals except carnivores, pinpointing this region as a likely interface of pathogen-driven adaptation. Importantly, several codons were detected under significant positive selection across multiple orders (Supplementary Table 2), including thirteen sites shared by at least three mammalian orders: one located in the RRM and twelve sites within the RING/DTC which exhibited important biochemical changes in hydrophobicity within and between orders (Figure 2d).

To assess the structural distribution of these adaptive changes, we modeled DTX3L structures from species of the three most strongly selected orders (bats, primates, and artiodactyls) using AlphaFold2 (Jumper et al., 2021). Positively selected sites were mainly located on the surfaces of RRM and RING-DTC domains, clustered in close spatial proximity, in the four mammalian orders analyzed (Figure 2e). In contrast, reported catalytic sites (707, 719 and 731, Dearlove et al., 2024) were highly conserved (Figure 2e), suggesting that substrate recognition and catalytic activity could be subject to shared selective pressures. In line with this, solvent accessibility analysis using FreeSASA (Mitternacht, 2016) showed that adaptive substitutions were enriched at exposed residues, with a significant over-representation in all species except rodents (Fisher’s exact test, odds ratio > 5; p < 0.005) (Supplementary Table 2). The signal was strongest for positively selected sites shared across mammals, which form a distinct surface cluster on the DTC domain, characteristic of substrate-binding interfaces and host–virus protein–protein coevolution (Figure 2f).

### Strong co-evolutionary dynamics between DTX3L and PARP9 have shaped the ubiquitin – ADP-ribose functional axis

Considering the functional and evolutionary interplay between DTX3L and its partners, PARP9 and PARP14, we next investigated whether PARP9 and PARP14 display parallel evolutionary trajectories with DTX3L. Applying the same evolutionary framework (aBSREL, BUSTED and site-specific analyses) to mammalian alignments, we showed that PARP9 and PARP14 showed similar evolutionary dynamics observed for DTX3L. First, as detected in DTX3L, bats, primates and artiodactyls harbored the highest proportion of significant positively selected codons (ranging 3.24 to 20.08%) (Supplementary Figure 3). Second, site-specific inference showed an enrichment of adaptive changes in the functional domains, particularly in the macrodomains in all mammals (Figure 3a). Importantly, a stretch of 19 and 9 positively selected sites located in macrodomains 1 and 2 of PARP9, respectively, were shared across mammalian orders (Figure 3a, b, Supplementary Table 2). This indicates convergent or shared selective pressures acting on these sites. This pattern was observed to a lesser extent in PARP14, in which three shared adaptive sites were identified in the MD3 and catalytic domains (CAT).

**Figure 3.**
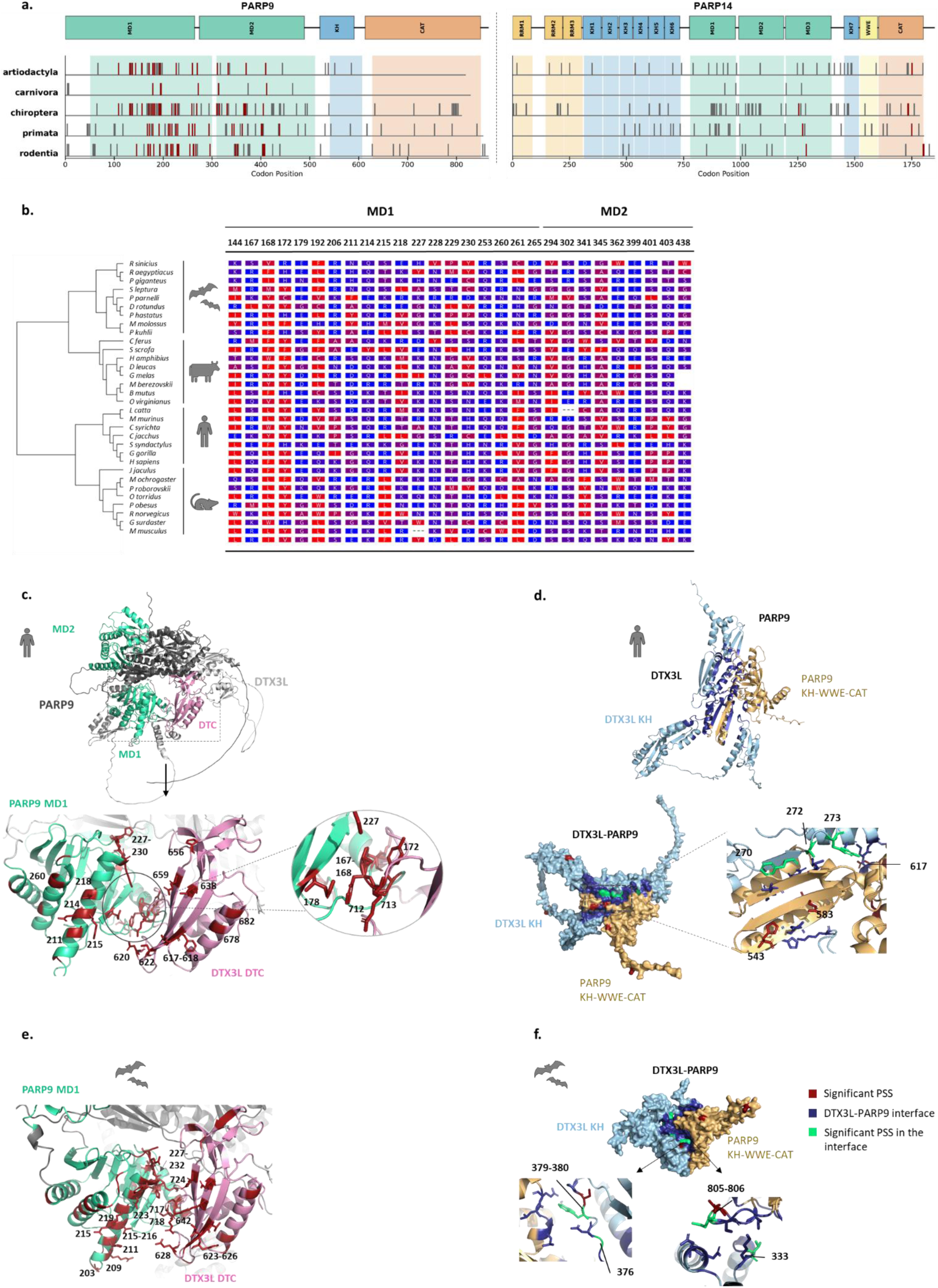
Coevolution of the DTX3L-PARP axis. **a)** Summary of BUSTED and models M0, M1, M2, M7, and M8 implemented in Bio++ and Codeml (PAML) with nested model comparisons (M1 vs. M2 and M7 vs. M8) assessed by likelihood ratio tests (LRTs) on PARP9 and PARP14. Colored boxes indicate models detecting a significant signal of positive selection and barplots indicate the proportion of all positively selected sites (Unique PSS) and Significant PSS (shared by at least two models). **b)** Significant positively selected sites mapped on the linear representation of PARP9 and PARP14 protein structures for each mammalian orders. **c)** Alignments of positively selected sites shared by at least three mammalian orders, highlighting amino acid changes in mammalian PARP9. The cladogram on the left shows the phylogenetic relationships among mammalian species. Codons with asterisk indicate the most rapidly evolving codons (BEB > 0.99 and detected by at least four methods). **d)** Predicted 3D docking structures of DTX3L-PARP9 complex from human using Colabfold (pLDDT>70), showing the spatial conformation of DTX3L RING-DTC and PARP9 macrodomain (left) and the enrichment of positively selected sites in both domains (right). **e)** Predicted 3D docking structures of DTX3L-PARP9 complex from human, with a close view of the positively selected sites in the interface (green). **f-g**) Similar analyses for the bat species *P. giganteus* as described for c-d.

Due to challenges in obtaining high-confidence protein structures for all three partners, we focused further analyses on the DTX3L-PARP9 complex, which forms a well-defined physical and functional unit. Combining site-inference analyses with protein structure docking, we uncovered a significant spatial concordance between the enrichment of positively selected sites in PARP9’s macrodomain 1 and the hotspot of genetic changes in the DTC domain of DTX3L (Figure 3c). More strikingly, several of these sites were structurally interacting (within 5 Å), and those at the interface correspond to shared positively selected sites across mammals in both proteins (Figure 3c). This suggests inter-residue co-evolution between the catalytic domains of DTX3L and PARP9. Among these sites, residues 620, 622, 659, 712, and 713 (DTX3L) and 167, 168, 172, 178, and 227 (PARP9, human sequence) were some of the fastest-evolving codons mapping directly to this interface, pinpointing these amino acids as critical candidates for interspecific functional differences.

In addition to the macrodomains, a number of adaptive sites also mapped in the KH and CAT domains of PARP9 and PARP14, which are known to mediate interactions with DTX3L (Kar et al., 2024) (Figure 3b, d). A similar pattern was reciprocally found in DTX3L, which also harbored positively selected sites in the KH domain that mediates interaction with PARP9 and PARP14. Modeling the 3D structure of the KH domain of DTX3L in complex with the KH–CAT region of PARP9, we showed that several of these positively selected sites mapped directly at, or in close proximity to the predicted interaction interface in primates (residues 270, 272, 274 in DTX3L and 543, 583, 617 in PARP9; Figure 3d). These patterns were found across the tested mammals, notably in bats, which showed the highest number of adaptive changes in all three protein partners (Figure 3e, f). Overall, these results indicate that selective pressures are unlikely to have acted on each partner individually, but have probably shaped the DTX3L-PARP9 interface as a whole component.

To test this hypothesis, we analyzed co evolutionary patterns between DTX3L and PARP9 using the MirrorTree method initially described in (Pazos & Valencia, 2001). Across all five mammalian orders, the two genes exhibited very high tree similarity (Pearson 0.946 ≤ r ≤ 0.982, highest p value p = 1.39e-67; Supplementary Table 2), indicating strongly correlated evolutionary distances between species. Topological consistency was highest in primates (Spearman ρ=0.972, p≈0), reflecting a very strong concordance in relative divergence rates between the two genes across lineages. Within each order, the minimum and maximum pairwise distances were highly similar between DTX3L and PARP9, indicating that both genes span nearly identical evolutionary depths. The larger maximum distances observed in rodents likely reflect a faster overall molecular evolutionary rate in this order rather than gene specific effects. Overall, these results point to co evolution signatures between DTX3L and PARP9 across all five mammalian orders, supporting long term functional coupling between the two genes. These results were further confirmed by the patterns of parallel episodic diversifying selection across the three genes, notably in bats and rodents, in which the same nodes and branches were inferred under significant positive selection (p-value < 0.05, Figure 4a, b), consistent with concordant adaptive responses and/or shared selective pressures.

**Figure 4.**
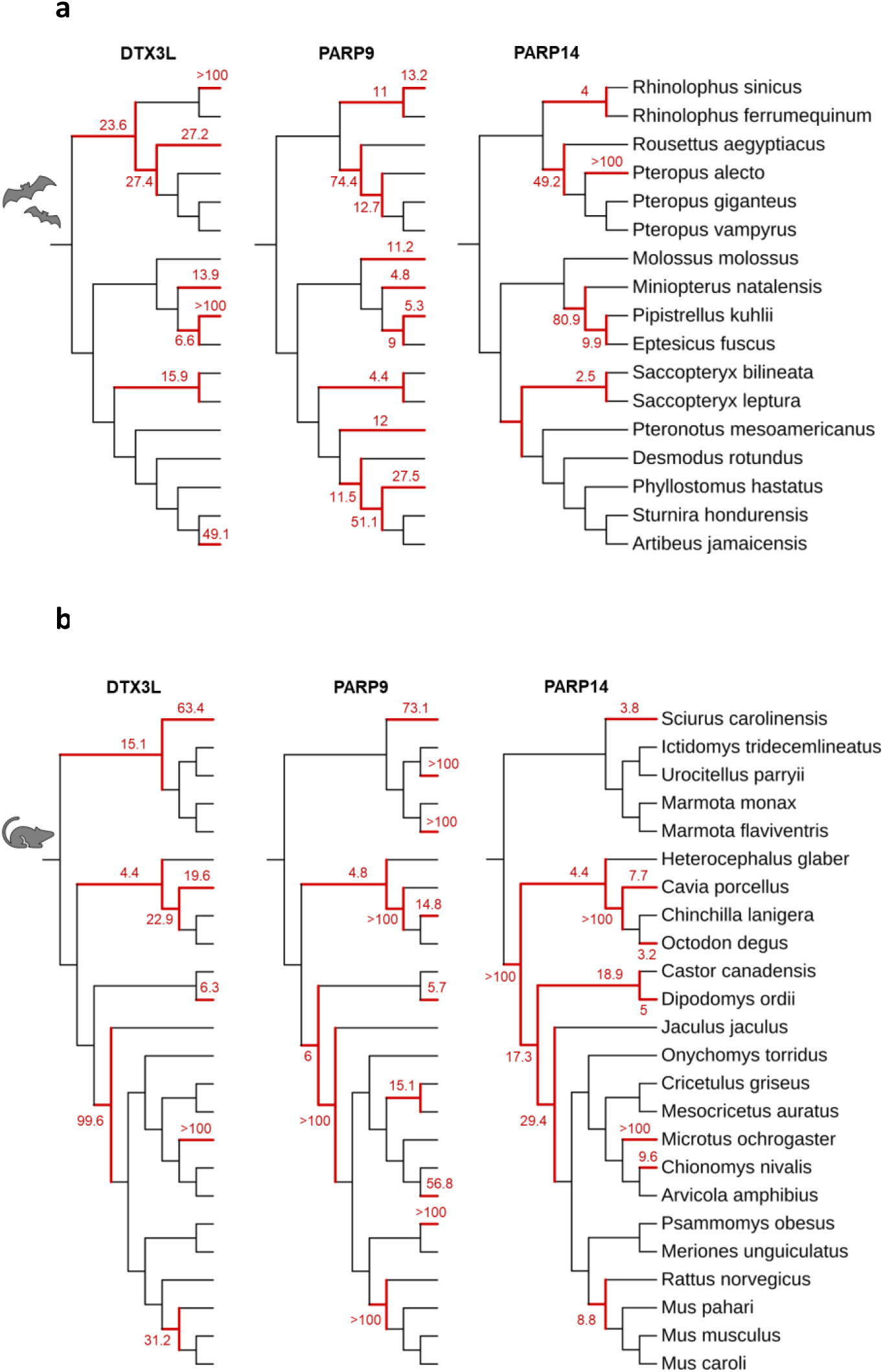
Concordant evolution across lineages. Phylogenetic trees of bats (left) and rodent (right) DTX3L, PARP9 and PARP14, showing similar branches and nodes under significant positive selection (p-value <0.05, in red).

### Short-timescale variation in the DTX3L–PARP complex suggests recent and ongoing evolution in mammals

Finally, we tested whether the signatures of recurrent adaptive evolution identified at the interspecific level (Figures 2, 3) also leave detectable footprints at shorter evolutionary timescales. To this end, we compiled non-synonymous Single-Nucleotide Polymorphisms (SNPs) from coding regions across datasets spanning a range of divergence levels, from populations within a single species to subspecies and closely related species within a genus. The sampled taxa encompass human (*Homo sapiens*), the *Chlorocebus* genus (comprising six lineages variously classified as species or subspecies; Svardal et al., 2017), dog (*Canis lupus familiaris*), cattle (*Bos taurus),* domestic sheep (*Ovis* genus), and domestic goat (*Capra* genus), as annotated in (Latrille et al., 2023). First, for each gene and dataset, we reconstructed amino-acid haplotype sequences and computed haplotype diversity (Hd), a measure of the probability that two randomly chosen haplotypes in a sample are distinct (Nei, 1987). This analysis revealed a clear asymmetry within the DTX family. While DTX2, DTX3, and DTX4 displayed low Hd values across all surveyed dataset (mean Hd = 0.13, 0.006, and 0.07, respectively), the three components of the DTX3L-PARP functional complex (DTX3L, PARP9, and PARP14) exhibited markedly elevated haplotypic diversity (mean Hd across all populations = 0.37, 0.41 and 0.59, respectively; Figure 5a). The concentration of standing variation specifically within the complex members corroborates the recurrent adaptive evolution signatures identified at the interspecific level (Figures 2, 3) and points to ongoing evolutionary dynamics within these genes. The elevated haplotypic diversity was particularly pronounced in *Chlorocebus* for all three complex genes, and shows variable distributions across the continent, with distinct haplotypic compositions observed between East African (*C. aethiops, C. p. hilgerti, C. cynosuros*), West African (*C. sabaeus*), Central African (*C. tantalus*), and Southern African (*C. p. pygerythrus*) populations (Figure 5b), confirming species-specific evolutionary trajectories.

**Figure 5.**
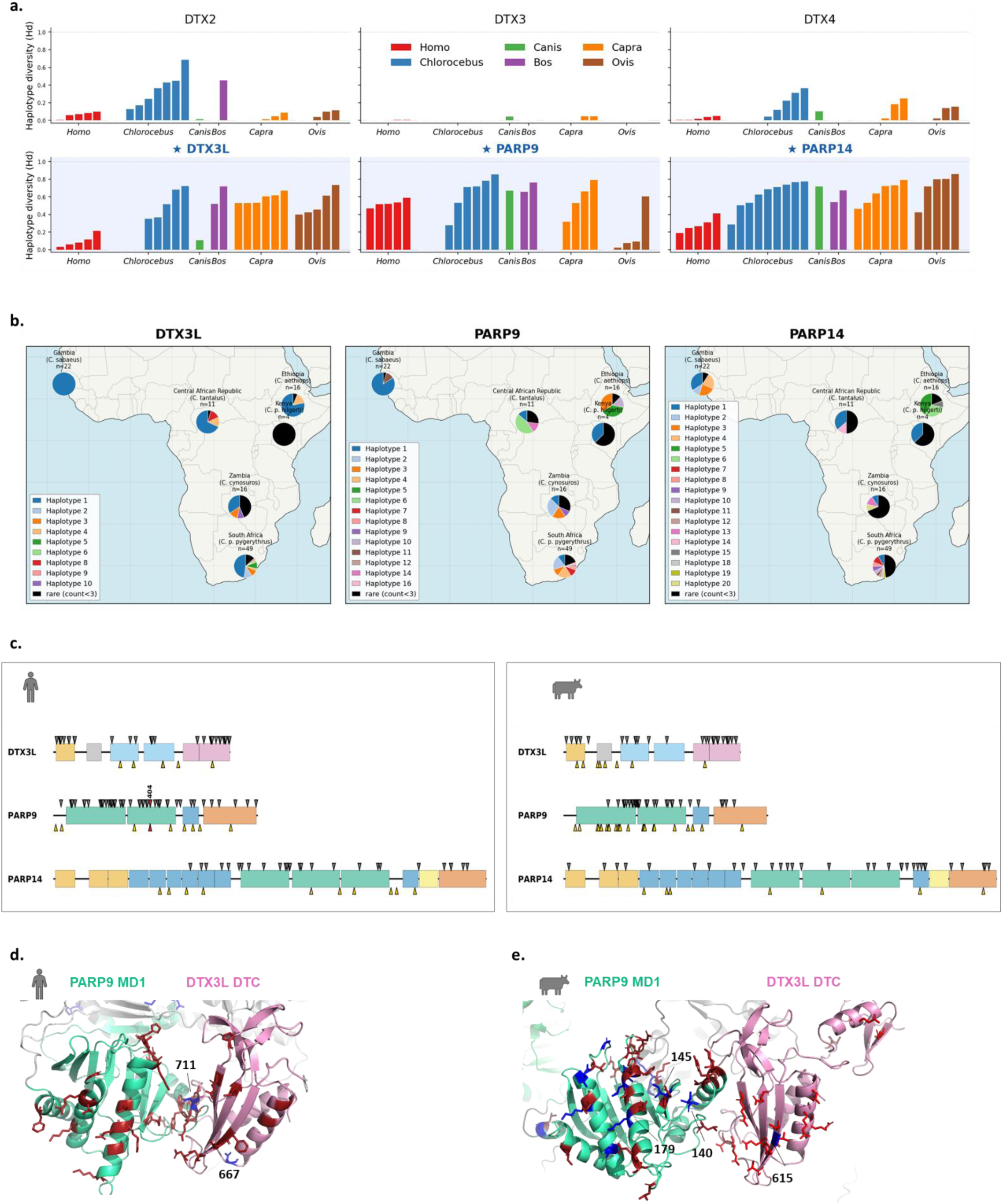
Ongoing variation in mammalian species of the DTX3L-PARP9-PARP14 complex. **a)** Haplotype diversity across the DTX-PARP gene family at shorter evolutionary timescales. Haplotype diversity (Hd = N/(N−1) × (1 − Σpᵢ²), where pᵢ is the frequency of haplotype i and N the number of samples; Nei, 1987) computed for each gene across datasets spanning different levels of divergence, from populations within a species to subspecies and closely related species within a genus. Each bar represents one dataset (population or lineage), colored and grouped by genus. The top row shows the non-complex DTX paralogs (DTX2, DTX3, DTX4) and the bottom row shows the three members of the DTX3L-PARP9-PARP14 functional complex (highlighted with a blue background). **b)** Geographic distribution of amino-acid haplotypes in *Chlorocebus* (Africa). Pie charts are positioned at the mean geographic coordinates of each sampled African lineage and show the relative frequency of each amino-acid haplotype. Each label indicates the *Chlorocebus* species or subspecies following the taxonomy of Svardal et al. (2017). Colours are gene-specific and are ranked by global frequency across the genus (Haplotype 1 = most frequent). Caribbean-origin populations (Barbados, Saint Kitts, Nevis) are excluded. Haplotypes observed in fewer than three individuals per dataset are pooled as ’rare’ (black). Panels from left to right: DTX3L, PARP9, PARP14. **c)** Distribution of positively selected sites (PSS) and non-synonymous single nucleotide polymorphisms (SNPs) across the coding regions of DTX3L, PARP9, and PARP14 for human and cattle. For each gene, PSS are shown along the top track in grey, and non-synonymous SNPs are shown along the bottom track in yellow. Sites that are both PSS and SNPs are highlighted in red, indicating positions where long-term adaptive evolution and ongoing intraspecific variation coincide. **d-e)** Predicted 3D docking structures of DTX3L-PARP9 complex from human **(d)** and cattle **(e)** using Colabfold (pLDDT>70), showing the spatial conformation of DTX3L RING-DTC and PARP9 macrodomain and the positions of the non-synonymous polymorphisms (blue) compared to the positively selected sites (red).

Projecting non-synonymous SNPs from coding regions onto gene-level alignments, we found species-specific genetic patterns at the protein level (Figure 5c). Despite this variability, the SNPs were enriched within the same domains identified at interspecific level (Figure 2, 3). Most non-synonymous SNPs mapped in the KH and RING-DTC domains of DTX3L, and the KH and macrodomains of PARP9 and PARP14 (Figure 5c, Supplementary Figure 4). Moreover, several SNPs clustered at the DTC/MD1 interface of the DTX3L–PARP9 complex in both human and cattle, including sites that overlap with or lie near residues under positive selection (Figure 5d). This overlap strongly suggests that these functional interfaces remain primary targets of ongoing evolutionary pressure.

### Viral pathogens as strong candidate drivers of DTX3L-PARPs axis diversification

To identify the viral candidates that could drive such evolutionary hallmarks, we interrogated the VirHostNet database (Guirimand et al., 2015). All three proteins of the DTX3L–PARPs complex were reported to be involved in interactions with various viral proteins (Figure 6), suggesting that it constitutes a recurrent interface in virus–host interactions. DTX3L is targeted by proteins encoded by Polyomaviridae and Papillomaviridae, including SV40 Large T antigen and HPV31 E4/E7. PARP9 displayed a broader viral interface, with interactors from human cytomegalovirus (HCMV), but shared the same interaction with SV40, suggesting this interaction targets the DTX3L-PARP9 dimer rather than individual genes. PARP9 and PARP14 show the largest number of reported interactors, including five proteins encoded by herpesviruses.

**Figure 6.**
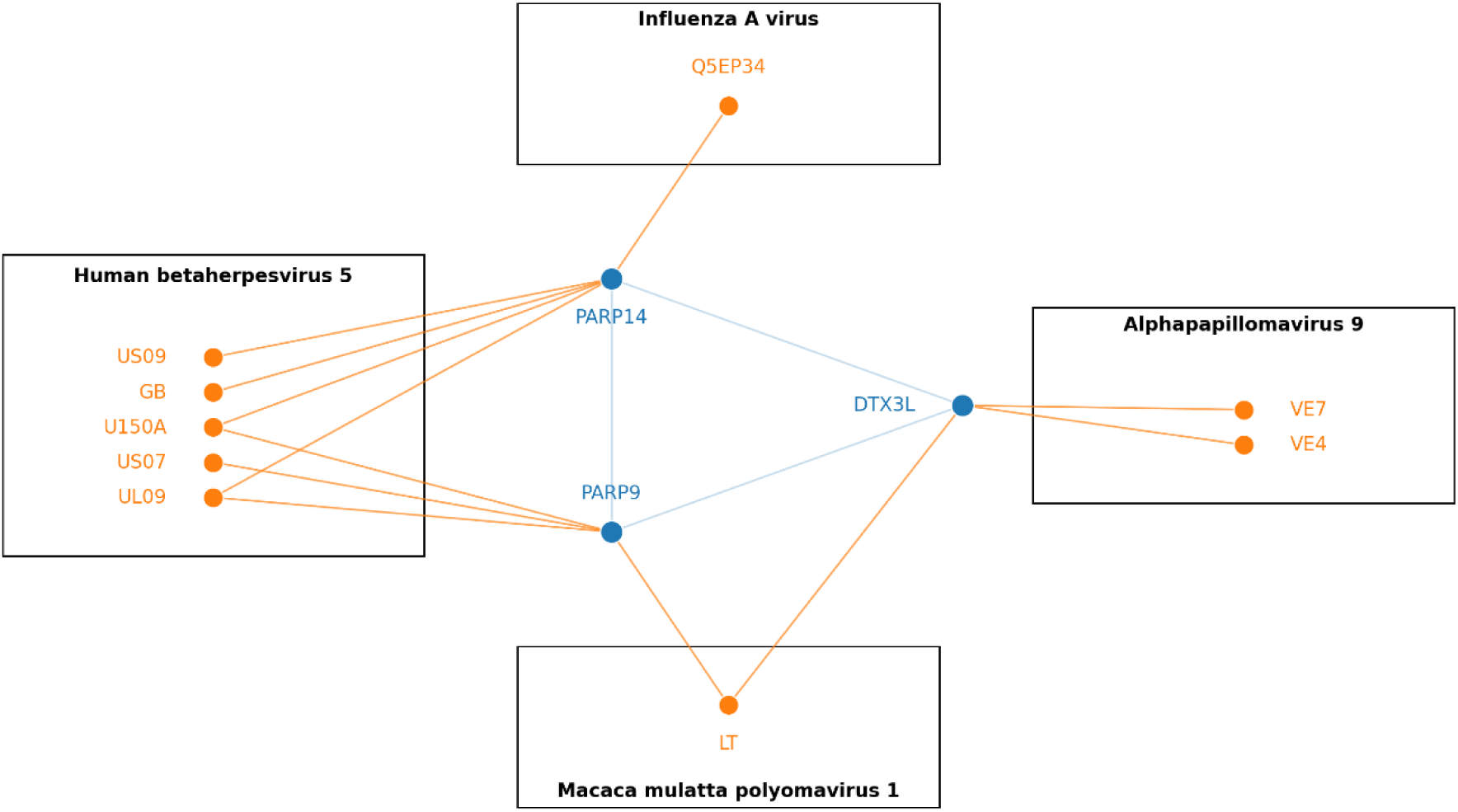
Reported interactions between viral proteins and the DTX3L-PARP9-PARP14 complex. Reported interactors were retrieved by querying the VirHostNet database for each protein individually as well as for the complex as a whole. In the resulting interaction network, blue vertices represent proteins that are part of the complex, and blue edges denote interactions among complex components. Orange vertices correspond to viral proteins, with orange edges indicating interactions between viral proteins and members of the host complex.

## DISCUSSION

While most phylogenomic studies of innate immunity tend to focus on specific proteins or gene families, here, we integrated the functional architecture of DTX3L–PARP9–PARP14 complex to decipher the evolutionary mechanisms underlying the diversification of ubiquitin – ADP-ribosylation axis. Our analyses reveal recurrent adaptive evolution of DTX3L and signatures of co-evolution across DTX3L-PARPs complex and across mammals, pointing to sustained evolutionary pressure on the ubiquitin–ADP-ribose regulatory axis. Notably, recurrent adaptive substitutions cluster within domains that are critical for catalytic activity, complex assembly, and structural stability, with significant hallmarks of concordant evolution between residues of the DTX3L-PARP9 complex. Moreover, multiple mammalian lineages exhibit signatures of convergent evolution across all three proteins, indicating that similar pressures have acted independently on the same pathway in different evolutionary contexts. Importantly, these genetic variations are still detectable over relatively recent evolutionary timescales, evidenced by the accumulation of non-synonymous polymorphisms within species. These findings suggest that the DTX3L–PARPs complex behave as an adaptive module, shaped by recurrent and presumably ongoing host–virus conflicts, highlighting its central role in vertebrate innate immunity.

All three components of the DTX3L-PARP complex are highly IFN-inducible in mammalian cells, with evidence of co-expression (Kar et al., 2024; Zhang et al., 2015). At the genomic level, DTX3L and PARP9 are arranged in a head-to-head orientation, and share a bidirectional IFN-γ–responsive promoter containing STAT and IRF binding sites (Juszczynski et al., 2006; Zhang et al., 2015). In human systems, perturbation of either gene alters the protein levels of its partner, revealing reciprocal regulation (Kar et al., 2024). Notably, PARP9 influences DTX3L at the post-transcriptional level, whereas DTX3L affects both PARP9 mRNA and protein abundance (Kar et al., 2024; Zhang et al., 2015). Extending previous genomic analyses beyond human, our findings indicate that this functional coupling is an ancient and conserved feature, at least at the mRNA level, across vertebrates, and emerged following the *Dipnoi* diversification. While the synteny between DTX3L and PARP9 was maintained since their emergence, intergenic distances vary widely, and inversely correlate with the strength of transcriptional coupling. This variation suggests that intergenic distance is a key parameter controlling co-expression strength. In this model, genomic linkage could allow coordinated expression, while genomic distance enables lineage-specific modulation of DTX3L-PARP9 co-expressions without disrupting functional coupling. In addition, the fact that PARP14 ranked among the top three co-expressed genes for DTX3L and PARP9 even in zebrafish, in which DTX3L is located in a distinct locus, indicates that the functional association among these partners predated the emergence of the DTX3L–PARP9 genomic configuration and reflects an ancient regulatory complex within vertebrates. While recent studies have highlighted the functional coupling of PARP14 with the DTX3L–PARP9 complex (Kar et al., 2024; Ribeiro et al., 2024), our results further suggest that this interaction may have arisen early through domain-level exchanges.

Beyond their functional interdependence, we show that DTX3L and PARP9 have evolved under concordant adaptive selection across mammals, suggesting that selection acts on this complex as a functional unit rather than as independent proteins. Reciprocal adaptation in regions required for DTX3L-PARP complex formation and function, coupled with convergent signatures of positive selection across multiple mammalian lineages, are consistent with classical signatures of co-evolving protein complexes under host-pathogen conflicts (Lovell & Robertson, 2010; Pazos & Valencia, 2008). These findings further align with the broader evolutionary principle that physical and/or functional interactions can constrain and link the evolutionary trajectories of protein partners (De Juan et al., 2013; Pazos & Valencia, 2008). Comparable dynamics, with concerted adaptive evolution, have been observed in antiviral pathways such as the OAS–RNase L and cGAS–STING axes, where positively selected codons cluster in functional and interaction domains (Chu et al., 2023). Although our data do not directly resolve co-evolution with PARP14, its functional coupling with the DTX3L–PARP9 complex, together with the cluster of adaptive changes within the same domains as PARP9, suggests a shared evolutionary trajectory. Altogether, these results support a model of co-evolution within the DTX3L–PARP complex, in which adaptive changes in one partner may drive functionally linked changes in others, strengthening their interdependence.

In this model, a plausible mechanism is that shared selective pressures act jointly on the catalytic domains of DTX3L and PARP9. This is supported by the accumulation of adaptive substitutions in the RING-DTC domain of DTX3L and the macrodomains of PARP9, which mediate ubiquitin transfer and ADP-ribose binding, respectively (Dearlove et al., 2024; Kar et al., 2024; C.-S. Yang et al., 2017). In the RING-DTC domain of DTX3L, catalytic residues required for ADPr ubiquitination (H707, E733, Y719) are strictly conserved (Dearlove et al., 2024; Zhu et al., 2024), whereas an adjacent cluster of rapidly evolving codons (616-712) delineates the DTC domain. This region likely defines a key host-pathogen interface, overlapping with sites critical for E2 enzyme interactions or substrate recognition (Dearlove et al., 2024; C.-S. Yang et al., 2017). Importantly, these adaptive residues structurally align with positively selected sites in macrodomain 1 of PARP9, suggesting a coordinated and conserved antagonistic mechanism in mammals. Such a model is consistent with known viral strategies to subvert ubiquitin and ADP-ribose signaling, such as encoding or hijacking RING-type E3 ligases (Zhang et al., 2018), or encoding macrodomain-containing proteins that bind or hydrolyze ADP-ribose (e.g. Fehr et al., 2016; Leung et al., 2018). While interactions between DTX3L and PARP9 have been primarily attributed to the KH domains (Chatrin et al., 2020; C.-S. Yang et al., 2017), these co-evolutionary signatures combined with the predicted spatial proximity (< 5 Å) between DTX3L RING-DTC and PARP9 macrodomain 1, point to a putative coordinated functional interface. Recent work on Zika virus further supports this model. Structural modeling predicts points of contact between the ZIKV envelope protein and both DTX3L and PARP9 (Gumpangseth et al., 2025). In our dataset, DTX3L S638, which is predicted to contact viral E protein, shows adaptive evolution in four mammalian orders, highlighting this site as a strong candidate of viral antagonism. Similarly, predicted ZIKV interacting residues in PARP9 map to macrodomain 1 and coincide with regions enriched in positive selection. Although these interactions remain to be experimentally validated, the convergence between predicted viral binding surfaces and adaptively evolving residues strongly supports a model in which viruses target the macrodomain–RING interface of the complex. In this context, while interactions between DTX3L and PARP9 have been primarily attributed to the KH domains, the co-evolutionary signatures and the predicted spatial proximity (< 5 Å) between the RING-DTC of DTX3L and the macrodomain 1 of PARP9, point to a putative coordinated functional interface.

A second, non-exclusive mechanism is the antagonism of the DTX3L–PARP9 interaction surface, which could impair or modulate complex formation between both partners. Although we detected fewer positively selected sites within the direct interface between DTX3L and PARP9, even limited amino acid changes at such interfaces can have significant functional effects, by altering binding affinity, specificity, or regulatory dynamics (De Juan et al., 2013; Pazos & Valencia, 2008). This suggests that selection may act not only on catalytic activity, but also on the stability and regulation of the complex itself.

Importantly, this signal is not restricted to deep evolutionary timescales. The presence of lineage-specific non-synonymous polymorphisms located near or overlapping with these sites suggests that the complex, and particularly this interface, remains under evolutionary pressure in contemporary populations. In line with this, the geographic structuration of haplotypes within *Chlorocebus* complex points to spatially heterogeneous selective pressures, possibly imposed by locally circulating viral communities, or with the complex demographic and taxonomic history of the genus *Chlorocebus* (Svardal et al., 2017). More broadly, the elevated haplotypic diversity observed for these genes, in contrast to other DTX family members suggests that selective pressures are specifically concentrated on this complex. Such past adaptations and ongoing standing variation in key binding surfaces may underlie differences in susceptibility to ancient and modern pathogens (Corona et al., 2018).

Beyond the interaction interface, our data indicate that adaptive selection also target distinct domain within DTX3L itself. Notably, we identified recurrent adaptive changes in the RRM domain across species, despite strong structural constraints imposed by its β-sheet nucleic acid–binding surface (Daubner et al., 2013). This pattern is noteworthy given that, within the DELTEX family, DTX3L and DTX3 are the only members harboring ssNA-ubiquitinating activity (Dearlove et al., 2024). The near-universal conservation of the DTX3L RRM further underscores its functional importance. The concentration of adaptive changes in this domain therefore points to an additional layer of functional conflict, independent of the DTX3L–PARP9 interface.

Such multi-layer functional conflict is consistent with known viral strategies. Several viral families, including SARS CoV 2 (Kar et al., 2024), paramyxoviruses (Huang et al., 2023) and picornaviruses (Zhang et al., 2015), are known to interfere with the complex. Notably, viral macrodomains from SARS CoV 2 have been shown to reverse PARP14 and DTX3L–PARP9 mediated modifications (Russo et al., 2021). Given the diversity of viral strategies to hijack, suppress, or redirect ubiquitin and ADP-ribosylation pathways (Fehr et al., 2020; Isaacson & Ploegh, 2009; Lant & Maluquer de Motes, 2021; Zhang et al., 2018), multiple other viruses could plausibly drive these evolutionary patterns. Herpesviruses are strong candidates as they encode multiple E3 ligases and ubiquitin-interacting proteins, and are known to extensively manipulate host ubiquitination pathways (Oswald et al., 2023; Soh et al., 2022), as supported by our VirHostNet analysis. Similarly, poxviruses encode multifunctional immune evasion proteins that broadly interfere with ubiquitin-dependent signaling (Lant & Maluquer de Motes, 2021), while RNA viruses such as flaviviruses are known to target ADP-ribosylation pathways (Malgras et al., 2021). In this context, the parallel accumulation of adaptive changes at the DTX3L–PARP9 interface and within the DTX3L RRM domain supports a model in which multiple functional surfaces of the complex are recurrently targeted by viral antagonists. Nonetheless, other pathogens beyond viruses may also contribute to the observed evolutionary patterns. In particular, intracellular bacteria are known to extensively manipulate host ubiquitination and ADP-ribosylation pathways, and could therefore impose selective pressures on the DTX3L–PARP9 axis (Roberts et al., 2023). In addition, while PARP9 alone exhibits limited catalytic activity and requires complex formation with DTX3L for full enzymatic and antiviral function (Zhang et al., 2015), DTX3L retains intrinsic E3 ubiquitin ligase activity and may participate in PARP9-independent pathways. Therefore, it cannot be excluded that additional cellular processes, such as regulation of ubiquitin-dependent signaling, DNA damage responses, or RNA-associated functions, have contributed to the evolutionary trajectory of DTX3L.

In conclusion, our findings indicate that through coordinated regulation and reciprocal adaptation of catalytic and binding surfaces, the DTX3L-PARP module is likely shaped by persistent pathogen pressure targeting the entire functional complex rather than individual components. We propose that viral antagonists targeting this system have evolved strategies to disrupt the ADPr–ubiquitin axis, thereby impairing ubiquitin tagging and the subsequent degradation of viral factors. These mechanisms may include: (i) antagonism of a catalytic domain in one partner with functional consequences for the other protein; (ii) simultaneous targeting of the catalytic domains of both proteins; and (iii) disruption of protein–protein interaction interfaces that stabilize complex formation. Our results further show the need to integrate both macro- and micro-evolutionary scales to decipher adaptive dynamics over evolutionary timescales. Further functional and mechanistic studies will be required to define the full extent and biological impact of these adaptive changes.

## MATERIAL AND METHODS

### Phylogenetic analyses in mammals

Protein sequences for each gene were retrieved from the NCBI databases for species representative of each order: B*os taurus, Canis lupus familiaris, Rhinolophus ferrumequinum, Homo sapiens, Mus musculus*. When multiple isoforms were available, the longest sequence was retained. Sequences identifiers are detailed in Supplementary Table 1.

Phylogenetic analyses of the seven genes of interest were conducted using DGINN commit a5928aa (Picard et al., 2020) across five mammalian orders. Orthologous groups were identified using the DGINN screening pipeline. Homologous nucleotide sequences were retrieved from the core_nt database using tblastn. Default DGINN parameters were applied for Primates and Carnivora (E-value ≤ 1 × 10⁻⁴; minimum identity ≥ 70%), whereas relaxed parameters were used for Artiodactyla, Chiroptera, and Rodentia (E-value = 0; minimum identity ≥ 60%). Sequences were first aligned using MAFFT v7.271 (Katoh & Standley, 2013) with default parameters, followed by codon-aware alignment using MACSE v2.07 (Ranwez et al., 2018). Maximum-likelihood phylogenetic trees were inferred using IQ-TREE v2.6.6 with automatic model selection (Minh et al., 2020). Following tree reconciliation with order-specific species trees, based on phylogenies from (Foley et al., 2023), using Treerecs v1.0 (Comte et al., 2020), orthologous groups were constituted, aligned and their phylogenetic trees reconstructed as previously described. Then, after manual curation, final codon-aware alignments and phylogenetic trees were generated in the same manner. Phylogenetic trees were visualized and edited using ITOL v7 (Letunic & Bork, 2024), available as a free web-server tool (https://itol.embl.de/).

### Positive selection analyses

Final codon-aware alignments and their corresponding phylogenetic trees were used to investigate signatures of positive selection at both site and branch levels. Specifically, BUSTED (Branch-site Unrestricted Statistical Test for Episodic Diversification) (Murrell et al., 2015), FUBAR (Fast Unconstrained Bayesian AppRoximation) (Murrell et al., 2013) and MEME (Mixed Effects Model of Evolution) (Murrell et al., 2012), implemented in HyPhy (Kosakovsky Pond et al., 2020), were applied to detect gene-wide and episodic site-specific selection, respectively. In parallel, site models (M0, M1, M2, M7, and M8) were tested using Bio++ (Guéguen et al., 2013) and the codeml program from PAML (Z. Yang, 2007) through ETE3 (v3.1.3, (Huerta-Cepas et al., 2016)). For both Bio++ and codeml, evidence of positive selection was evaluated using likelihood ratio tests (LRTs) comparing nested models (M1 vs. M2 and M7 vs. M8). Sites were considered statistically significant if they met at least one of the following criteria: (i) posterior probability > 0.95 under the Bayes Empirical Bayes (BEB) framework in codeml, (ii) posterior probability > 0.95 in Bio++ or FUBAR, or (iii) p < 0.05 in MEME. To minimize false positives, only sites identified by at least two independent methods were retained as candidates for positive selection.

Branch-specific selection analyses were conducted using HYPHY aBSREL (adaptive Branch-Site REL) v2.5.72 (Smith et al., 2015) across vertebrates, and across each of the five mammalian orders. To enable comparison of branch-specific selection among genes within the same order, analyses were restricted to species shared among genes within each order, and the corresponding pruned alignments and trees were used for aBSREL analyses.

Enrichment of positively selected sites was statistically inferred using the Kruskal–Wallis (KW) test. When the KW test was significant (p < 0.05), post hoc pairwise comparisons were performed using Wilcoxon rank-sum tests, with p-values adjusted for multiple comparisons using the Benjamini–Holm correction.

### DTX3L duplication and genomic synteny analysis

To retrace the phylogenetic history of the DELTEX family, we created a custom database of 126 NCBI RefSeq genomes covering 21 classes of Metazoa (Supplementary table 2). For each order, we selected two species representing at least two distinct families, prioritizing chromosome-level assemblies. Homologs of the genes of interest were identified using tblastn searches with human protein sequences as queries (Supplementary Table 2) under relaxed thresholds (e-value ≤ 0.01; ≥ 30% identity). Following manual curation, sequences were aligned in subgroups of approximately 200 sequences using MAFFT (E-INS-i model). Sub-alignments were subsequently merged using a global pair alignment strategy. Phylogenetic relationships within the family were inferred using IQ-TREE under the Q.MAMMAL+R8 substitution model, with branch support assessed using ultra-fast bootstrap (1000 replicates) (Hoang et al., 2018) and approximate likelihood ratio test (aLRT, 1000 replicates) (Anisimova & Gascuel, 2006).

The synteny between DTX3L and PARP9 was analyzed across metazoan species using Ensembl and RefSeq genome annotations. We restricted these analyses to species with chromosome-level or high-quality scaffold assemblies. Gene orientation, adjacency, and intergenic distances were derived from genomic coordinates of annotated transcripts, with a focus on the longest protein-coding isoforms. Shared untranslated regions (UTRs) were defined as direct overlaps between annotated UTR exons of DTX3L and PARP9. To control for annotation artifacts, UTR configurations were compared across independent annotation databases (ENSEMBL, NCBI Refseq). Evolutionary conservation of the intergenic region was assessed using UCSC multi-species genome alignments generated with MultiZ, providing additional support for the authenticity of observed structural variation. As a control, the genomic organization of genes flanking the DTX3L–PARP9 locus was analyzed using the same criteria as for the focal gene pair. For each species, gene identity, orientation, and intergenic distances of neighboring genes were compared across taxa to assess whether the variability observed at the DTX3L–PARP9 locus reflects locus-specific constraints rather than genome-wide structural variation.

### Co-expression analyses

Top 100 co-expressed genes for each DTX3L, PARP9 and PARP14 were extracted from COXPRESdb (https://coxpresdb.jp/top_search/#CoExSearch) (Obayashi et al., 2008) for eight available species: human, mouse, rat, monkey, dog, cat, chicken, and zebrafish (Supplementary Table 1). Gene ranks, based on Z-scores, were normalized to percentile values to enable cross-species comparisons of co-expression patterns. Pairwise and triple overlaps among the top co-expressed genes were quantified, and the statistical significance of enrichment was assessed using hypergeometric tests based on species-specific genome sizes. The correlation between genomic distance and co-expression Z-score was assessed in species exhibiting conserved DTX3L–PARP9 synteny using Spearman’s rank correlation test implemented in R. Results were visualized using the ggplot2 package in R.

### Protein structure modeling and protein-protein complex docking

To investigate the spatial distribution of adaptive sites, we characterized them on the 3D structure from the three representative species for orders harboring the highest number of adaptive sites (bats, primates, and artiodactyls). For primates, the predicted DTX3L structure from *Homo sapiens* was obtained from AlphaFold Protein Structure Database (<u>alphafold.ebi.ac.uk</u>). For bats and artiodactyls, the 3D structure of DTX3L from respectively *Pteropus giganteus* and *Bos mutus* were generated using AlphaFold2 (Jumper et al., 2021) via the Neurosnap platform (https://neurosnap.ai/). The top ranked models were selected according to pLDDT confidence scores (>70). Visualization and mapping of positively selected sites were done using PyMOL v3.1.3.1. Relative Solvent accessibility (RSA) of each residue was computed using FreeSASA (Mitternacht, 2016). Residues were categorized as solvent-exposed or buried according to RSA thresholds, and the enrichment of positively selected sites among exposed residues was statistically evaluated using Fisher’s exact test.

To assess whether positively selected sites located in interaction regions were spatially associated with interface residues between DTX3L and PARP9, we generated a structural model of the KH domain of DTX3L in complex with the KH-CAT domains of PARP9 for human and bats, using *P giganteus* as a reference species, and ColabFold (Mirdita et al., 2022). The top-ranked model based on interface predicted TM-score (ipTM, mean ipTM > 0.85) was selected for downstream analyses and visualization in PyMOL v3.1.3.1.

Interface residues were identified through electrostatic interface analysis combined with spatial proximity criteria (residues within 5 Å across chains, with high-confidence scores) and positively selected sites were mapped onto the predicted complex structure.

### Co-evolution analyses

Protein co evolution was estimated using the MirrorTree approach to quantify the similarity between phylogenetic trees of protein families as a proxy for functional association (Pazos & Valencia, 2001). Briefly, alignments of DTX3L and PARP9 obtained previously for each order were converted to amino-acids, their evolutionary distance matrices were computed using protdist (version 3.5c) and the Pearson and Spearman coefficients were calculated as a measure of co-evolution between the two genes.

### Virus – host protein interaction analyses

To identify viral proteins potentially interacting with the DTX3L–PARP9–PARP14 complex in mammals, we queried the publicly available VirHostNet database (https://virhostnet.prabi.fr/virhostevol/) (Guirimand et al., 2015). Viral interacting proteins were retrieved independently for each host gene (DTX3L, PARP9 and PARP14) as well as for the complex as a whole, based on reported host–virus protein–protein interaction data.

### Single-nucleotide polymorphisms analyses

To assess ongoing microevolutionary dynamics in DTX3L, we analyzed single-nucleotide polymorphisms (SNPs) in the DTX3L-PARP9-PARP14 axis across six genera covering three mammalian orders: *Bos taurus, Ovis,* and *Capra* (Artiodactyla); *Homo sapiens* and *Chlorocebus* (Primates); and *Canis lupus familiaris* (Carnivora). The polymorphism datasets were obtained as detailed in (Latrille et al., 2023). SNPs were mapped onto the same reference gene sequences used for interspecific analyses, enabling direct comparison of their distribution patterns across evolutionary scales.

### Data and code availability

DGINN is available for download at https://github.com/leapicard/DGINN. All alignments, phylogenetic trees and output for the different analyses are available at https://doi.org/10.5281/zenodo.19691367. Original data for single-nucleotide analyses are available at https://doi.org/10.5281/zenodo.7107233.

## ACKNOWLEDGMENTS

We thank the Zoonoses Virales team (MIVEGEC Lab), and the Cell Signal Unit members (OIST) for thoughtful discussions and feedback. We are grateful for the help and support provided by the Scientific Computing and Data Analysis section of Core Facilities at OIST. We would also like to thank Dr L. Gueguen (LBBE) for his help and for the maintenance of DGINN.

## FUNDING

SJ was supported by the CNRS and the French Agence Nationale de la Recherche (ANR), under grant ANR-25-CE35-6757-01. LP was supported by JSPS KAKENHI grant number JP25K18819. TL was supported by Université de Lausanne and the Swiss National Science Foundation (315230_219757).

## AUTHOR CONTRIBUTIONS

Conceptualization: SJ. Methodology: LP, SJ. Formal analysis: LP, EE, TL, SJ. Visualization: LP, EE, TL. Funding: SJ. Writing – original draft: LP, SJ. Review: all authors.

## COMPETING INTERESTS

The authors declare that they have no competing interests.

## Notes

https://doi.org/10.5281/zenodo.19691367

### Competing Interest Statement

The authors have declared no competing interest.

## REFERENCES

Agrata, R., & Komander, D. (2025). Ubiquitin—A structural perspective. Molecular Cell, 85(2), 323–346. 10.1016/j.molcel.2024.12.015

Anisimova, M., & Gascuel, O. (2006). Approximate Likelihood-Ratio Test for Branches: A Fast, Accurate, and Powerful Alternative. Systematic Biology, 55(4), 539–552. 10.1080/10635150600755453

Barbour, H., Nkwe, N. S., Estavoyer, B., Messmer, C., Gushul-Leclaire, M., Villot, R., Uriarte, M., Boulay, K., Hlayhel, S., Farhat, B., Milot, E., Mallette, F. A., Daou, S., & Affar, E. B. (2023). An inventory of crosstalk between ubiquitination and other post-translational modifications in orchestrating cellular processes. iScience, 26(5). 10.1016/j.isci.2023.106276

Chatrin, C., Gabrielsen, M., Buetow, L., Nakasone, M. A., Ahmed, S. F., Sumpton, D., Sibbet, G. J., Smith, B. O., & Huang, D. T. (2020). Structural insights into ADP-ribosylation of ubiquitin by Deltex family E3 ubiquitin ligases. Science Advances, 6(38), eabc0418. 10.1126/sciadv.abc0418

Chatrin, C., Zhu, K., & Ahel, I. (2025). The rise of ADP-ribose–ubiquitin. Nature Structural & Molecular Biology, 32(9), 1582–1585. 10.1038/s41594-025-01651-0

Chu, L., Gong, Z., Wang, W., & Han, G.-Z. (2023). Origin of the OAS–RNase L innate immune pathway before the rise of jawed vertebrates via molecular tinkering. Proceedings of the National Academy of Sciences, 120(31), e2304687120. 10.1073/pnas.2304687120

Comte, N., Morel, B., Hasic, D., Guéguen, L., Boussau, B., Daubin, V., Penel, S., Scornavacca, C., Gouy, M., Stamatakis, A., Tannier, E., & Parsons, D. P. (2020). Treerecs: An integrated phylogenetic tool, from sequences to reconciliations. *Bioinformatics*, btaa615. 10.1093/bioinformatics/btaa615

Corona, E., Wang, L., Ko, D., & Patel, C. J. (2018). Systematic detection of positive selection in the human-pathogen interactome and lasting effects on infectious disease susceptibility. PLOS ONE, 13(5), e0196676. 10.1371/journal.pone.0196676

Dasovich, M., & Leung, A. K. L. (2023). PARPs and ADP-ribosylation: Deciphering the complexity with molecular tools. Molecular Cell, 83(10), 1552–1572. 10.1016/j.molcel.2023.04.009

Daubner, G. M., Cléry, A., & Allain, F. H.-T. (2013). RRM–RNA recognition: NMR or crystallography…and new findings. *Current Opinion in Structural Biology*, Folding and Binding / Protein-Nucleic Acid Interactions, 23(1), 100–108. 10.1016/j.sbi.2012.11.006

Daugherty, M. D., Young, J. M., Kerns, J. A., & Malik, H. S. (2014). Rapid Evolution of PARP Genes Suggests a Broad Role for ADP-Ribosylation in Host-Virus Conflicts. PLoS Genetics, 10(5), e1004403. 10.1371/journal.pgen.1004403

De Juan, D., Pazos, F., & Valencia, A. (2013). Emerging methods in protein co-evolution. Nature Reviews Genetics, 14(4), 249–261. 10.1038/nrg3414

Dearlove, E. L., Chatrin, C., Buetow, L., Ahmed, S. F., Schmidt, T., Bushell, M., Smith, B. O., & Huang, D. T. (2024). DTX3L ubiquitin ligase ubiquitinates single-stranded nucleic acids. eLife, 13, RP98070. 10.7554/eLife.98070

Du, Q., Miao, Y., He, W., & Zheng, H. (2023). ADP-Ribosylation in Antiviral Innate Immune Response. Pathogens (Basel, Switzerland), 12(2), 303. 10.3390/pathogens12020303

Fehr, A. R., Channappanavar, R., Jankevicius, G., Fett, C., Zhao, J., Athmer, J., Meyerholz, D. K., Ahel, I., & Perlman, S. (2016). The Conserved Coronavirus Macrodomain Promotes Virulence and Suppresses the Innate Immune Response during Severe Acute Respiratory Syndrome Coronavirus Infection. mBio, 7(6), 10.1128/mbio.01721-16. 10.1128/mbio.01721-16

Fehr, A. R., Singh, S. A., Kerr, C. M., Mukai, S., Higashi, H., & Aikawa, M. (2020). The impact of PARPs and ADP-ribosylation on inflammation and host-pathogen interactions. Genes & Development, 34(5–6), 341–359. 10.1101/gad.334425.119

Foley, N. M., Mason, V. C., Harris, A. J., Bredemeyer, K. R., Damas, J., Lewin, H. A., Eizirik, E., Gatesy, J., Karlsson, E. K., Lindblad-Toh, K., Zoonomia Consortium, Springer, M. S., & Murphy, W. J. (2023). A genomic timescale for placental mammal evolution. Science, 380(6643), eabl8189. 10.1126/science.abl8189

Guéguen, L., Gaillard, S., Boussau, B., Gouy, M., Groussin, M., Rochette, N. C., Bigot, T., Fournier, D., Pouyet, F., Cahais, V., Bernard, A., Scornavacca, C., Nabholz, B., Haudry, A., Dachary, L., Galtier, N., Belkhir, K., & Dutheil, J. Y. (2013). Bio++: Efficient Extensible Libraries and Tools for Computational Molecular Evolution. Molecular Biology and Evolution, 30(8), 1745–1750. 10.1093/molbev/mst097

Guirimand, T., Delmotte, S., & Navratil, V. (2015). VirHostNet 2.0: Surfing on the web of virus/host molecular interactions data. Nucleic Acids Research, 43(D1), D583–D587. 10.1093/nar/gku1121

Gumpangseth, N., Supungul, D., Villarroel, P. M. S., Songhong, T., Oabsuwan, K., Yainoy, S., Hamel, R., Missé, D., Monteil, A., Meewan, I., Saetear, P., & Wichit, S. (2025). E3 Ligase DTX3L and PARP9 Cooperatively Restrict Zika Virus by Modulating TLR/RLR-Mediated Innate Immunity and Targeting the Viral Envelope Protein (SSRN Scholarly Paper No. 5806047). Social Science Research Network. 10.2139/ssrn.5806047

Hoang, D. T., Chernomor, O., von Haeseler, A., Minh, B. Q., & Vinh, L. S. (2018). UFBoot2: Improving the Ultrafast Bootstrap Approximation. Molecular Biology and Evolution, 35(2), 518–522. 10.1093/molbev/msx281

Hsieh, H.-C., Ling, L.-L., & Wang, Y.-C. (2026). Post-translational modifications of immune checkpoints: Molecular mechanisms, tumor microenvironment remodeling, and therapeutic implications. Journal of Biomedical Science, 33(1), 3. 10.1186/s12929-025-01202-1

Huang, J., Chen, Z., Ye, Y., Shao, Y., Zhu, P., Li, X., Ma, Y., Xu, F., Zhou, J., Wu, M., Gao, X., Yang, Y., Zhang, J., & Hao, C. (2023). DTX3L Enhances Type I Interferon Antiviral Response by Promoting the Ubiquitination and Phosphorylation of TBK1. Journal of Virology, 97(6), e00687–23. 10.1128/jvi.00687-23

Huerta-Cepas, J., Serra, F., & Bork, P. (2016). ETE 3: Reconstruction, Analysis, and Visualization of Phylogenomic Data. Molecular Biology and Evolution, 33(6), 1635–1638. 10.1093/molbev/msw046

Isaacson, M. K., & Ploegh, H. L. (2009). Ubiquitination, Ubiquitin-like Modifiers, and Deubiquitination in Viral Infection. Cell Host & Microbe, 5(6), 559–570. 10.1016/j.chom.2009.05.012

Jacquet, S., El Filali, A., Etienne, L., & Pontier, D. (2025). Extensive adaptive changes in bat interferon pathway reveal specific molecular functions at the forefront of host–virus coevolution. Evolutionary Biology. 10.1101/2025.09.17.676773

Jumper, J., Evans, R., Pritzel, A., Green, T., Figurnov, M., Ronneberger, O., Tunyasuvunakool, K., Bates, R., Žídek, A., Potapenko, A., Bridgland, A., Meyer, C., Kohl, S. A. A., Ballard, A. J., Cowie, A., Romera-Paredes, B., Nikolov, S., Jain, R., Adler, J., … Hassabis, D. (2021). Highly accurate protein structure prediction with AlphaFold. Nature, 596(7873), 583–589. 10.1038/s41586-021-03819-2

Juszczynski, P., Kutok, J. L., Li, C., Mitra, J., Aguiar, R. C. T., & Shipp, M. A. (2006). BAL1 and BBAP Are Regulated by a Gamma Interferon-Responsive Bidirectional Promoter and Are Overexpressed in Diffuse Large B-Cell Lymphomas with a Prominent Inflammatory Infiltrate. Molecular and Cellular Biology, 26(14), 5348–5359. 10.1128/MCB.02351-05

Kar, P., Chatrin, C., Đukić, N., Suyari, O., Schuller, M., Zhu, K., Prokhorova, E., Bigot, N., Baretić, D., Ahel, J., Elsborg, J. D., Nielsen, M. L., Clausen, T., Huet, S., Niepel, M., Sanyal, S., Ahel, D., Smith, R., & Ahel, I. (2024). PARP14 and PARP9/DTX3L regulate interferon-induced ADP-ribosylation. The EMBO Journal, 43(14), 2929–2953. 10.1038/s44318-024-00126-0

Katoh, K., & Standley, D. M. (2013). MAFFT Multiple Sequence Alignment Software Version 7: Improvements in Performance and Usability. Molecular Biology and Evolution, 30(4), 772–780. 10.1093/molbev/mst010

Kosakovsky Pond, S. L., Poon, A. F. Y., Velazquez, R., Weaver, S., Hepler, N. L., Murrell, B., Shank, S. D., Magalis, B. R., Bouvier, D., Nekrutenko, A., Wisotsky, S., Spielman, S. J., Frost, S. D. W., & Muse, S. V. (2020). HyPhy 2.5—A Customizable Platform for Evolutionary Hypothesis Testing Using Phylogenies. Molecular Biology and Evolution, 37(1), 295–299. 10.1093/molbev/msz197

Lant, S., & Maluquer de Motes, C. (2021). Poxvirus Interactions with the Host Ubiquitin System. Pathogens, 10(8), 1034. 10.3390/pathogens10081034

Latrille, T., Rodrigue, N., & Lartillot, N. (2023). Genes and sites under adaptation at the phylogenetic scale also exhibit adaptation at the population-genetic scale. Proceedings of the National Academy of Sciences, 120(11), e2214977120. 10.1073/pnas.2214977120

Letunic, I., & Bork, P. (2024). Interactive Tree of Life (iTOL) v6: Recent updates to the phylogenetic tree display and annotation tool. Nucleic Acids Research, 52(W1), W78–W82. 10.1093/nar/gkae268

Leung, A. K. L., McPherson, R. L., & Griffin, D. E. (2018). Macrodomain ADP-ribosylhydrolase and the pathogenesis of infectious diseases. PLOS Pathogens, 14(3), e1006864. 10.1371/journal.ppat.1006864

Li, Z., Luo, A., & Xie, B. (2023). The Complex Network of ADP-Ribosylation and DNA Repair: Emerging Insights and Implications for Cancer Therapy. International Journal of Molecular Sciences, 24(19), 15028. 10.3390/ijms241915028

Lovell, S. C., & Robertson, D. L. (2010). An Integrated View of Molecular Coevolution in Protein–Protein Interactions. Molecular Biology and Evolution, 27(11), 2567–2575. 10.1093/molbev/msq144

Malgras, M., Garcia, M., Jousselin, C., Bodet, C., & Lévêque, N. (2021). The Antiviral Activities of Poly-ADP-Ribose Polymerases. Viruses, 13(4), 582. 10.3390/v13040582

Millán-Zambrano, G., Burton, A., Bannister, A. J., & Schneider, R. (2022). Histone post-translational modifications—Cause and consequence of genome function. Nature Reviews Genetics, 23(9), 563–580. 10.1038/s41576-022-00468-7

Minh, B. Q., Schmidt, H. A., Chernomor, O., Schrempf, D., Woodhams, M. D., von Haeseler, A., & Lanfear, R. (2020). IQ-TREE 2: New Models and Efficient Methods for Phylogenetic Inference in the Genomic Era. Molecular Biology and Evolution, 37(5), 1530–1534. 10.1093/molbev/msaa015

Mirdita, M., Schütze, K., Moriwaki, Y., Heo, L., Ovchinnikov, S., & Steinegger, M. (2022). ColabFold: Making protein folding accessible to all. Nature Methods, 19(6), 679–682. 10.1038/s41592-022-01488-1

Mitternacht, S. (2016). FreeSASA: An open source C library for solvent accessible surface area calculations. F1000Research, 5, 189. 10.12688/f1000research.7931.1

Murrell, B., Moola, S., Mabona, A., Weighill, T., Sheward, D., Kosakovsky Pond, S. L., & Scheffler, K. (2013). FUBAR: A Fast, Unconstrained Bayesian AppRoximation for Inferring Selection. Molecular Biology and Evolution, 30(5), 1196–1205. 10.1093/molbev/mst030

Murrell, B., Weaver, S., Smith, M. D., Wertheim, J. O., Murrell, S., Aylward, A., Eren, K., Pollner, T., Martin, D. P., Smith, D. M., Scheffler, K., & Kosakovsky Pond, S. L. (2015). Gene-Wide Identification of Episodic Selection. Molecular Biology and Evolution, 32(5), 1365–1371. 10.1093/molbev/msv035

Murrell, B., Wertheim, J. O., Moola, S., Weighill, T., Scheffler, K., & Kosakovsky Pond, S. L. (2012). Detecting Individual Sites Subject to Episodic Diversifying Selection. PLoS Genetics, 8(7), e1002764. 10.1371/journal.pgen.1002764

Nag, J., Patel, J., & Tripathi, S. (2023). Ubiquitin-Mediated Regulation of Autophagy During Viral Infection. Current Clinical Microbiology Reports, 10(1), 1–8. 10.1007/s40588-022-00186-y

Nei, M. (1987). Molecular evolutionary genetics. Columbia University Press.

Obayashi, T., Hayashi, S., Shibaoka, M., Saeki, M., Ohta, H., & Kinoshita, K. (2008). COXPRESdb: A database of coexpressed gene networks in mammals. Nucleic Acids Research, 36(suppl_1), D77–D82. 10.1093/nar/gkm840

Oswald, J., Constantine, M., Adegbuyi, A., Omorogbe, E., Dellomo, A. J., & Ehrlich, E. S. (2023). E3 Ubiquitin Ligases in Gammaherpesviruses and HIV: A Review of Virus Adaptation and Exploitation. Viruses, 15(9), 1935. 10.3390/v15091935

Palazzo, L., Mikoč, A., & Ahel, I. (2017). ADP -ribosylation: New facets of an ancient modification. The FEBS Journal, 284(18), 2932–2946. 10.1111/febs.14078

Pazos, F., & Valencia, A. (2001). Similarity of phylogenetic trees as indicator of protein–protein interaction. *Protein Engineering*, Design and Selection, 14(9), 609–614. 10.1093/protein/14.9.609

Pazos, F., & Valencia, A. (2008). Protein co-evolution, co-adaptation and interactions. The EMBO Journal, 27(20), 2648–2655. 10.1038/emboj.2008.189

Pereira-Leal, J. B., Levy, E. D., & Teichmann, S. A. (2006). The origins and evolution of functional modules: Lessons from protein complexes. Philosophical Transactions of the Royal Society B: Biological Sciences, 361(1467), 507–517. 10.1098/rstb.2005.1807

Picard, L., Ganivet, Q., Allatif, O., Cimarelli, A., Guéguen, L., & Etienne, L. (2020). DGINN, an automated and highly-flexible pipeline for the detection of genetic innovations on protein-coding genes. Nucleic Acids Research, 48(18), e103–e103. 10.1093/nar/gkaa680

Ranwez, V., Douzery, E. J. P., Cambon, C., Chantret, N., & Delsuc, F. (2018). MACSE v2: Toolkit for the Alignment of Coding Sequences Accounting for Frameshifts and Stop Codons. Molecular Biology and Evolution, 35(10), 2582–2584. 10.1093/molbev/msy159

Ribeiro, V. C., Russo, L. C., & Hoch, N. C. (2024). PARP14 is regulated by the PARP9/DTX3L complex and promotes interferon γ-induced ADP-ribosylation. The EMBO Journal, 43(14), 2908–2928. 10.1038/s44318-024-00125-1

Roberts, C. G., Franklin, T. G., & Pruneda, J. N. (2023). Ubiquitin-targeted bacterial effectors: Rule breakers of the ubiquitin system. The EMBO Journal, 42(18), EMBJ2023114318. 10.15252/embj.2023114318

Russo, L. C., Tomasin, R., Matos, I. A., Manucci, A. C., Sowa, S. T., Dale, K., Caldecott, K. W., Lehtiö, L., Schechtman, D., Meotti, F. C., Bruni-Cardoso, A., & Hoch, N. C. (2021). The SARS-CoV-2 Nsp3 macrodomain reverses PARP9/DTX3L-dependent ADP-ribosylation induced by interferon signaling. The Journal of Biological Chemistry, 297(3), 101041. 10.1016/j.jbc.2021.101041

Saleh, H., Liloglou, T., Rigden, D. J., Parsons, J. L., & Grundy, G. J. (2024). KH-like Domains in PARP9/DTX3L and PARP14 Coordinate Protein–Protein Interactions to Promote Cancer Cell Survival. Journal of Molecular Biology, 436(4), 168434. 10.1016/j.jmb.2023.168434

Scalia, P., Williams, S. J., Suma, A., & Carnevale, V. (2023). The DTX Protein Family: An Emerging Set of E3 Ubiquitin Ligases in Cancer. Cells, 12(13), 1680. 10.3390/cells12131680

Smith, M. D., Wertheim, J. O., Weaver, S., Murrell, B., Scheffler, K., & Kosakovsky Pond, S. L. (2015). Less Is More: An Adaptive Branch-Site Random Effects Model for Efficient Detection of Episodic Diversifying Selection. Molecular Biology and Evolution, 32(5), 1342–1353. 10.1093/molbev/msv022

Soh, S.-M., Kim, Y.-J., Kim, H.-H., & Lee, H.-R. (2022). Modulation of Ubiquitin Signaling in Innate Immune Response by Herpesviruses. International Journal of Molecular Sciences, 23(1), 492. 10.3390/ijms23010492

Svardal, H., Jasinska, A. J., Apetrei, C., Coppola, G., Huang, Y., Schmitt, C. A., Jacquelin, B., Ramensky, V., Müller-Trutwin, M., Antonio, M., Weinstock, G., Grobler, J. P., Dewar, K., Wilson, R. K., Turner, T. R., Warren, W. C., Freimer, N. B., & Nordborg, M. (2017). Ancient hybridization and strong adaptation to viruses across African vervet monkey populations. Nature Genetics, 49(12), 1705–1713. 10.1038/ng.3980

Takeyama, K., Aguiar, R. C. T., Gu, L., He, C., Freeman, G. J., Kutok, J. L., Aster, J. C., & Shipp, M. A. (2003). The BAL-binding Protein BBAP and Related Deltex Family Members Exhibit Ubiquitin-Protein Isopeptide Ligase Activity *. Journal of Biological Chemistry, 278(24), 21930–21937. 10.1074/jbc.M301157200

Thirunavukkarasu, S., Ahmed, M., Rosa, B. A., Boothby, M., Cho, S. H., Rangel-Moreno, J., Mbandi, S. K., Schreiber, V., Gupta, A., Zuniga, J., Mitreva, M., Kaushal, D., Scriba, T. J., & Khader, S. A. (2023). Poly(ADP-ribose) polymerase 9 mediates early protection against *Mycobacterium tuberculosis* infection by regulating type I IFN production. The Journal of Clinical Investigation, 133(12). 10.1172/JCI158630

Vozandychova, V., Stojkova, P., Hercik, K., Rehulka, P., & Stulik, J. (2021). The Ubiquitination System within Bacterial Host–Pathogen Interactions. Microorganisms, 9(3), 638. 10.3390/microorganisms9030638

Wang, L., Sun, X., He, J., & Liu, Z. (2021). Functions and Molecular Mechanisms of Deltex Family Ubiquitin E3 Ligases in Development and Disease. Frontiers in Cell and Developmental Biology, 9. 10.3389/fcell.2021.706997

Yan, Q., Ding, J., Khan, S. J., Lawton, L. N., & Shipp, M. A. (2023). DTX3L E3 ligase targets p53 for degradation at poly ADP-ribose polymerase-associated DNA damage sites. iScience, 26(4), 106444. 10.1016/j.isci.2023.106444

Yang, C.-S., Jividen, K., Spencer, A., Dworak, N., Ni, L., Oostdyk, L. T., Chatterjee, M., Kuśmider, B., Reon, B., Parlak, M., Gorbunova, V., Abbas, T., Jeffery, E., Sherman, N. E., & Paschal, B. M. (2017). Ubiquitin Modification by the E3 Ligase/ADP-Ribosyltransferase Dtx3L/Parp9. Molecular Cell, 66(4), 503–516.e5. 10.1016/j.molcel.2017.04.028

Yang, Z. (2007). PAML 4: Phylogenetic Analysis by Maximum Likelihood. Molecular Biology and Evolution, 24(8), 1586–1591. 10.1093/molbev/msm088

Ye, Q., Ma, J., Wang, Z., Li, L., Liu, T., Wang, B., Zhu, L., Lei, Y., Xu, S., Wang, K., Jian, Y., Ma, B., Fan, Y., Liu, J., Gao, Y., Huang, H., & Li, L. (2024). DTX3L-mediated TIRR nuclear export and degradation regulates DNA repair pathway choice and PARP inhibitor sensitivity. Nature Communications, 15(1), 10596. 10.1038/s41467-024-54978-5

Zhang, Y., Li, L.-F., Munir, M., & Qiu, H.-J. (2018). RING-Domain E3 Ligase-Mediated Host–Virus Interactions: Orchestrating Immune Responses by the Host and Antagonizing Immune Defense by Viruses. Frontiers in Immunology, 9. 10.3389/fimmu.2018.01083

Zhang, Y., Mao, D., Roswit, W. T., Jin, X., Patel, A. C., Patel, D. A., Agapov, E., Wang, Z., Tidwell, R. M., Atkinson, J. J., Huang, G., McCarthy, R., Yu, J., Yun, N. E., Paessler, S., Lawson, T. G., Omattage, N. S., Brett, T. J., & Holtzman, M. J. (2015). PARP9-DTX3L ubiquitin ligase targets host histone H2BJ and viral 3C protease to enhance interferon signaling and control viral infection. Nature Immunology, 16(12), 1215–1227. 10.1038/ni.3279

Zhu, K., Chatrin, C., Smith, R., Ahel, D., & Ahel, I. (2025). Interplay between ubiquitination and ADP-ribosylation and the case of dual modification ADPr-Ub. Essays in Biochemistry, 69(04), 267–279. 10.1042/EBC20253040

Zhu, K., Suskiewicz, M. J., Chatrin, C., Strømland, Ø., Dorsey, B. W., Aucagne, V., Ahel, D., & Ahel, I. (2024). DELTEX E3 ligases ubiquitylate ADP-ribosyl modification on nucleic acids. Nucleic Acids Research, 52(2), 801–815. 10.1093/nar/gkad1119.

