## Supplementary Figures for "DTX3L–PARP9 co-evolve as a single adaptive unit linking ubiquitination and ADP-ribosylation in antiviral immunity"

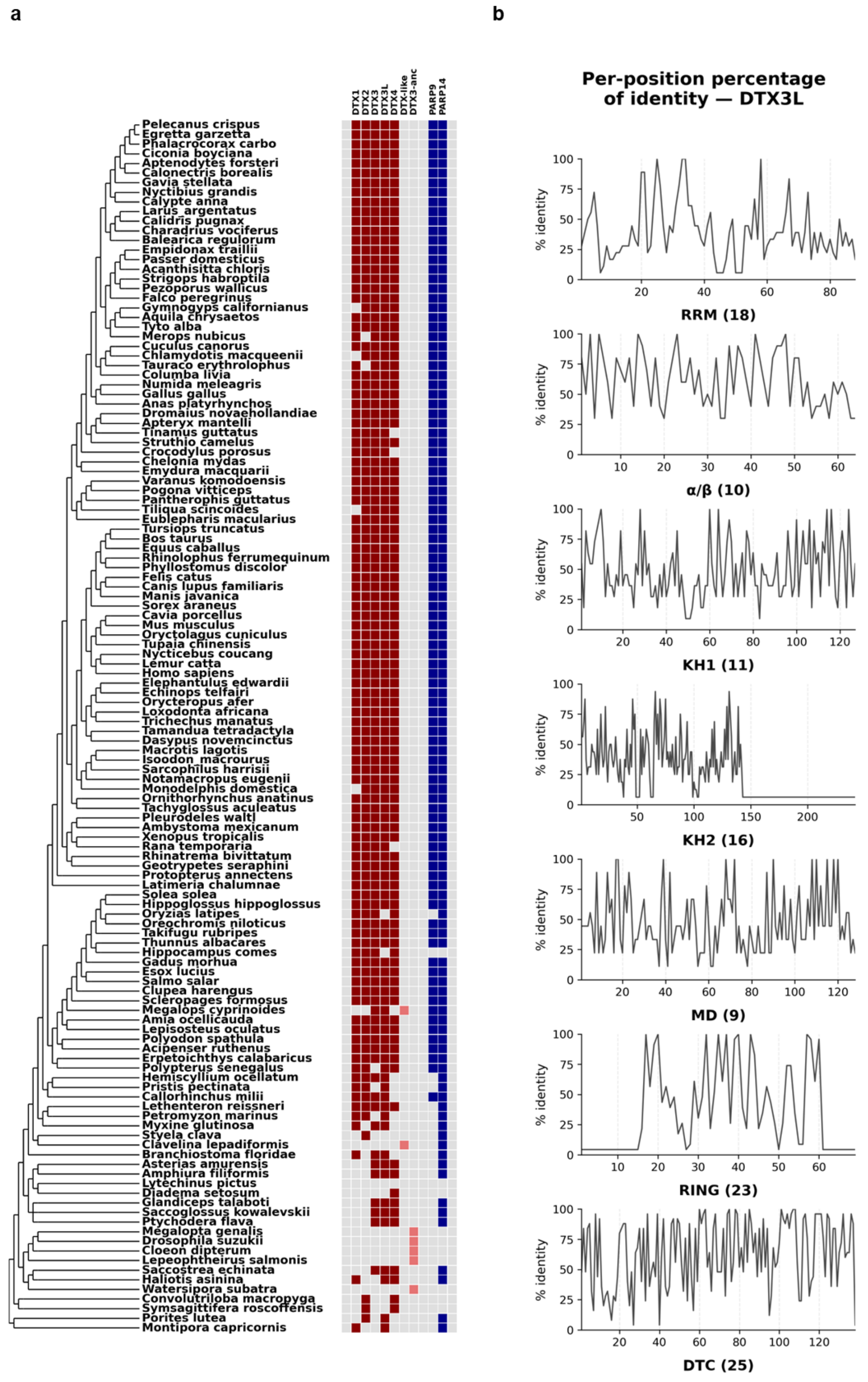


***Supplementary Figure 1.*** **a)** Phylogenetic profiling of Deltex and PARP genes in 126 metazoan species. Filled cells indicate the presence of an ortholog, while empty cells denote absence. Deltex family proteins are shown in red (lighter red for inferred ancestral Deltex proteins) and PARP proteins in blue. **b)** Per-position percentage of amino-acid identity across predicted DTX3L domains, highlighting regions of high conservation versus rapid divergence.

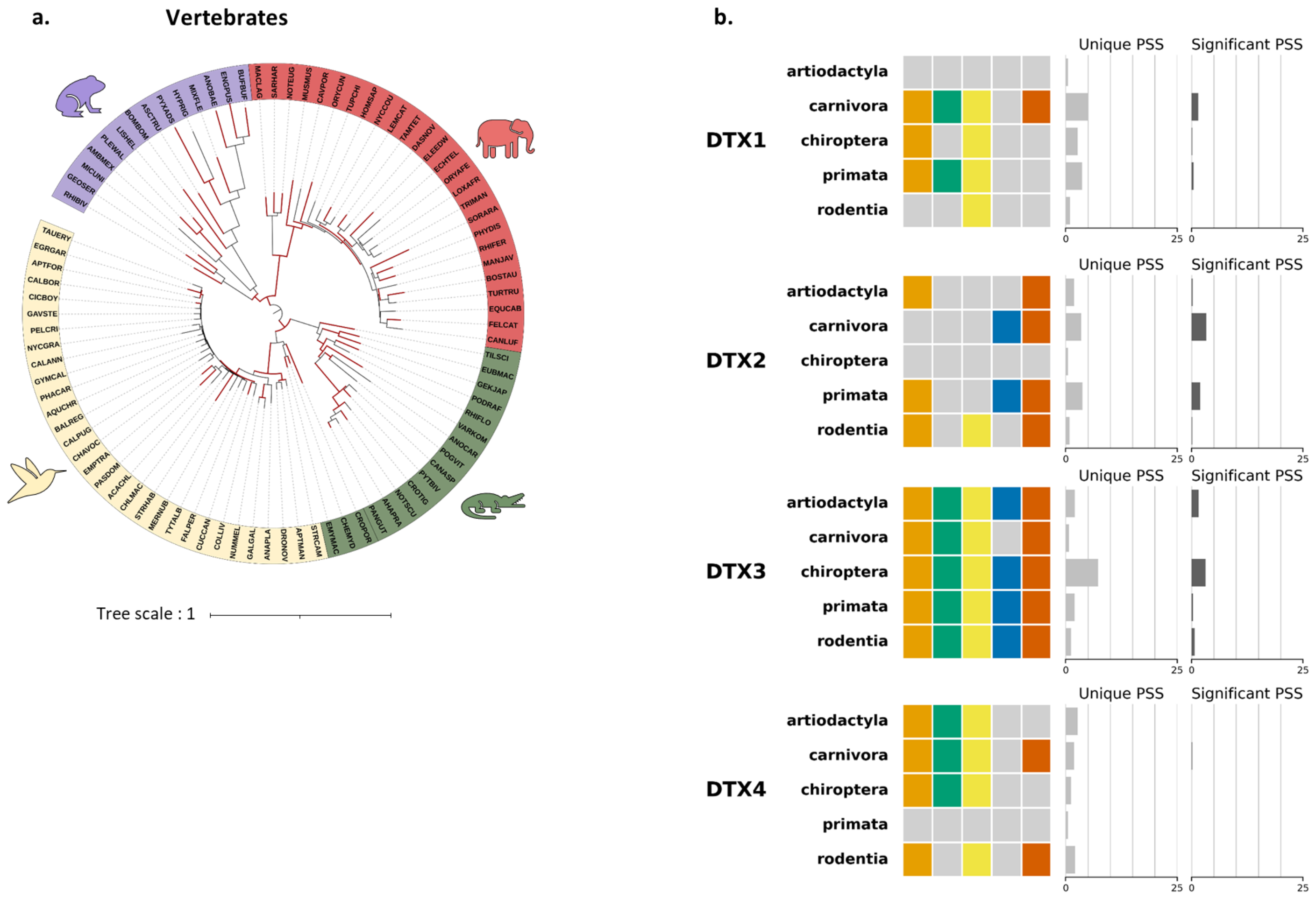


**Supplementary Figure 2. a)** Adaptive Branch-Site Random Effects Likelihood (aBSREL) analysis of DTX3L phylogeny across vertebrates. Red branches indicate lineages with significant evidence of positive selection (p < 0.05). **b)** Summary of BUSTED and models M0, M1, M2, M7, and M8 implemented in Bio++ and Codeml (PAML) assessing positive selection in the DTX family members across mammals. Colored boxes indicate models detecting a significant signal of positive selection and barplots indicate the proportion of all positively selected sites (Unique PSS) and Significant PSS (shared by at least two models).

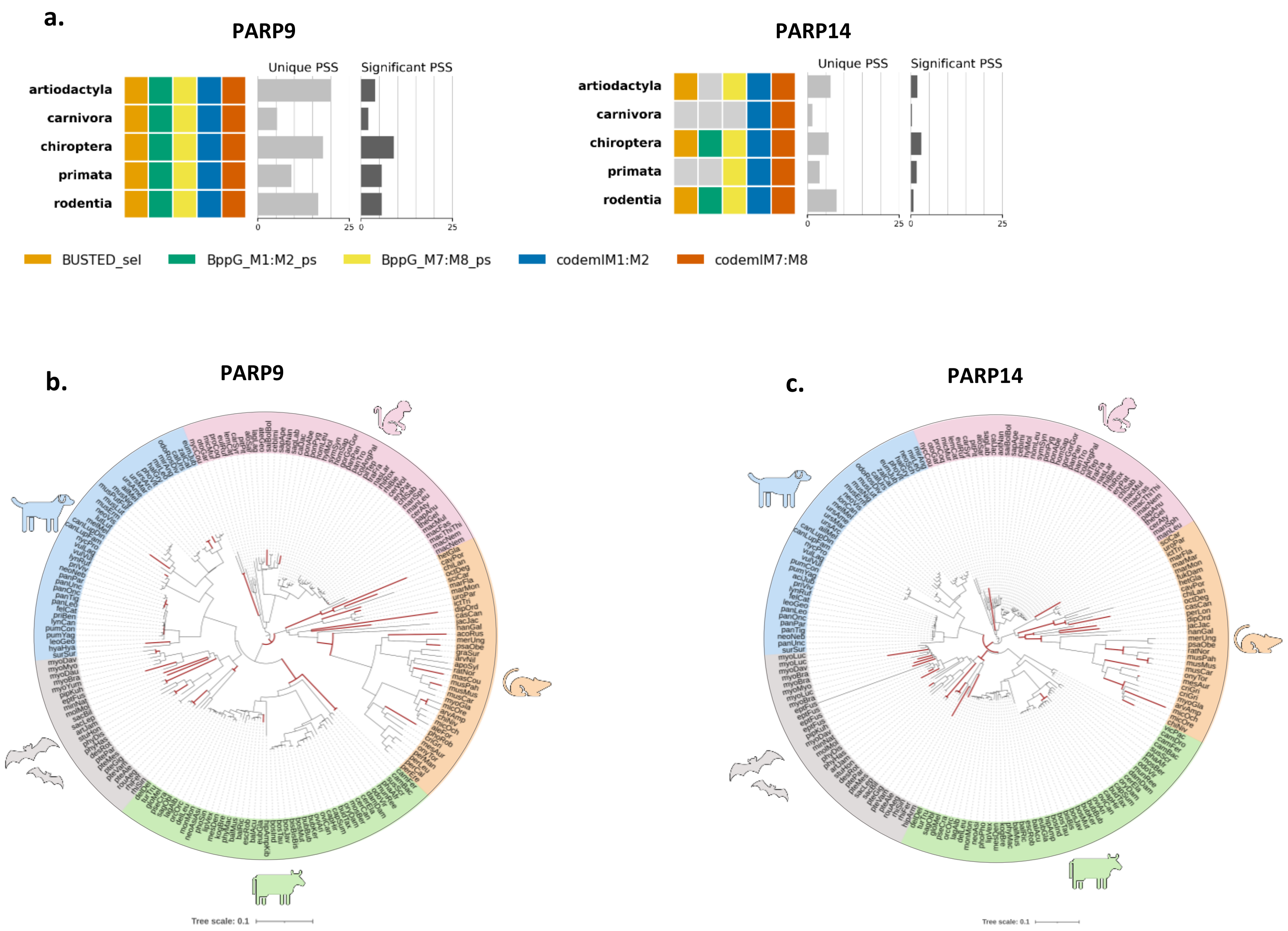


***Supplementary Figure 3.*** **Episodic positive selection of PARP9 and PARP14 mirrors the evolutionary pattern observed for DTX3L across mammals. a)** Summary of BUSTED and models M0, M1, M2, M7, and M8 implemented in Bio++ and Codeml (PAML) with nested model comparisons (M1 vs. M2 and M7 vs. M8) assessed by likelihood ratio tests (LRTs) on PARP9 and PARP14. Adaptive Branch-Site Random Effects Likelihood (aBSREL) analysis of PARP9 **(b)** and PARP14 **(c)** phylogenies across major mammalian lineages (primates, rodents, artiodactyls, bats, and carnivores). Red branches indicate lineages with significant evidence of positive selection ($p < 0.05$).

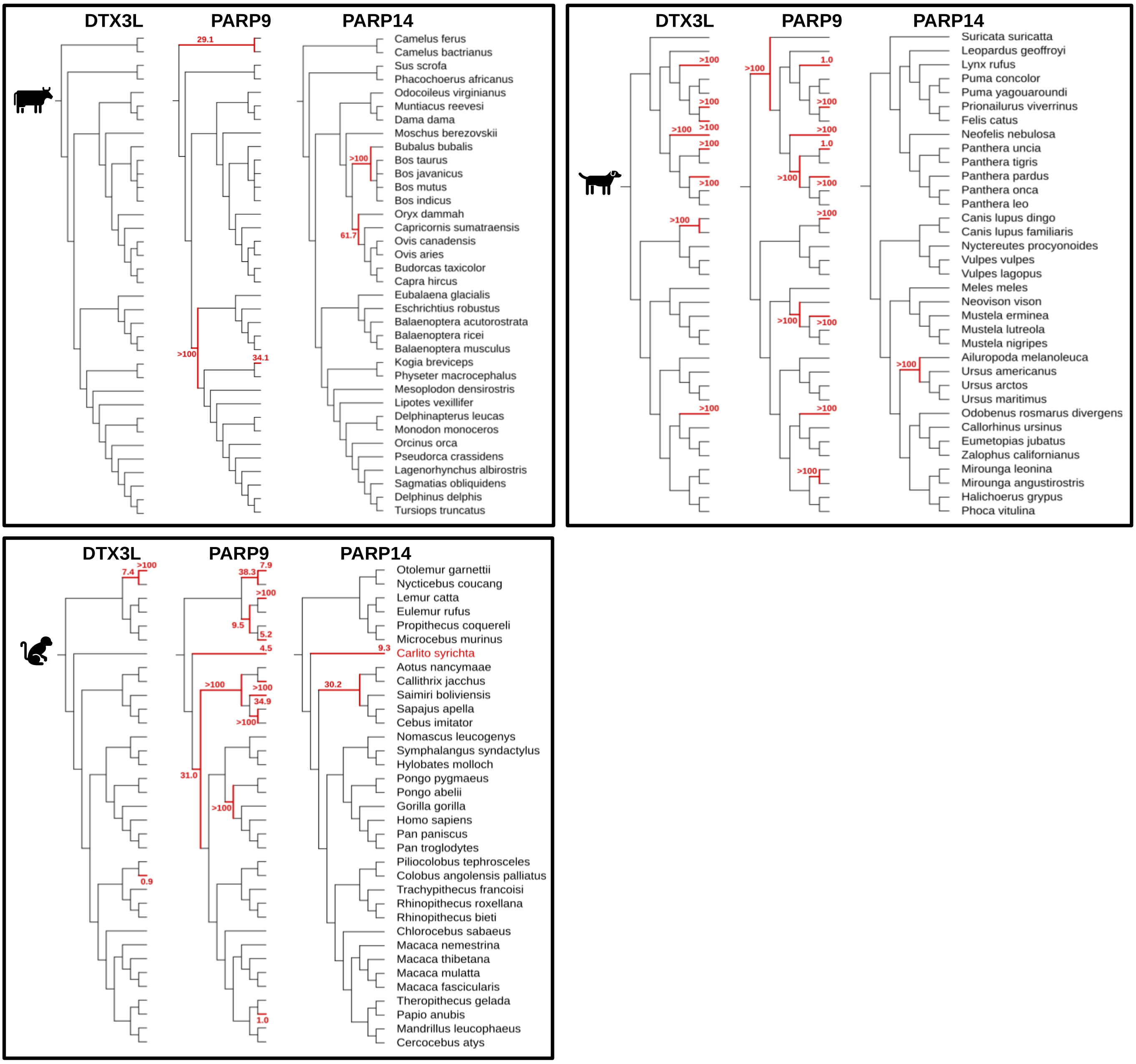


***Supplementary Figure 4.*** Cladograms of artiodactyl, carnivore, and primate lineages for DTX3L, PARP9, and PARP14. Episodic positive selection was evaluated using aBSREL, on the gene tree. Red branches show lineages under significant positive selection ($p < 0.05$).

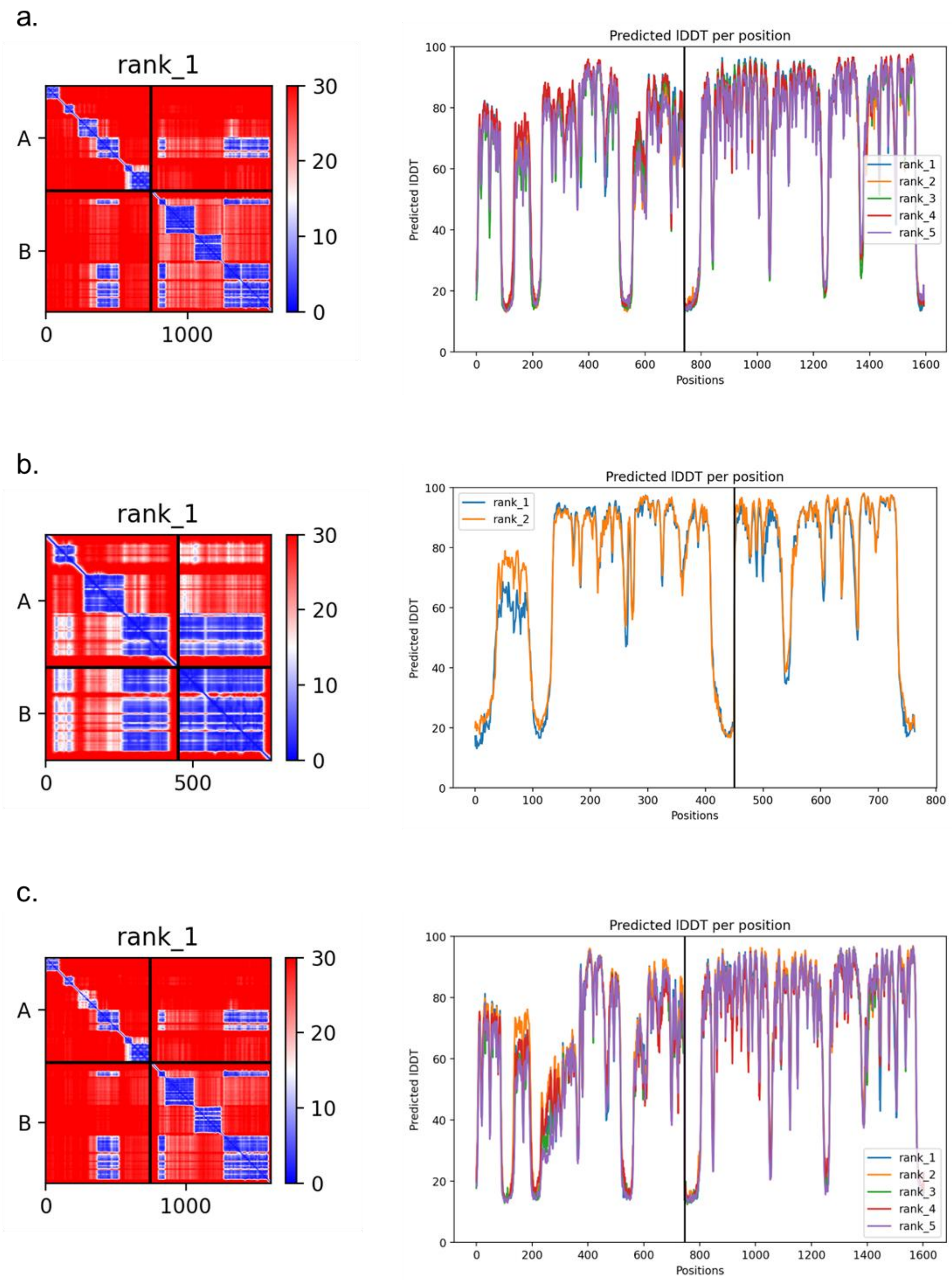


***Figure 5.*** Confidence analysis of protein–protein docking predictions generated with ColabFold for human (a), cattle (b), and bats (c). Left: predicted aligned error (PAE) plots for DTX3L (1–740) and PARP9 (741–1601). Right: predicted pLDDT scores per residue across ranked models.
